# A new computational method for membrane compressibility: Bilayer mechanical thickness revisited

**DOI:** 10.1101/360792

**Authors:** M. Doktorova, M.V. LeVine, G. Khelashvili, H. Weinstein

**Affiliations:** Tri-Institutional PhD Program in Computational Biology and Medicine, Weill Cornell Medical College, 1300 York Ave, New York NY 10065, USA; Department of Physiology and Biophysics, Weill Cornell Medicine, 1300 York Ave, New York NY 10065, USA; The HRH Prince Alwaleed Bin Talal Bin Abdulaziz Alsaud Institute for Computational Biomedicine, 13th Floor, Weill Greenberg Center, 1305 York Ave, New York NY 10065, USA

## Abstract

Because lipid bilayers can bend and stretch in ways similar to thin elastic sheets, physical models of bilayer deformation have utilized mechanical constants such as the moduli for bending rigidity (*κ_C_*) and area compressibility (*K_A_*). However, the use of these models to quantify the energetics of membrane deformation associated with protein-membrane interactions and the membrane response to stress is often hampered by the shortage of experimental data suitable for the estimation of the mechanical constants of various lipid mixtures. While computational tools such as Molecular Dynamics (MD) simulations can provide alternative means to estimate *K_A_* values, current approaches suffer significant technical limitations. Here, we present a novel computational framework that allows for a direct estimation of *K_A_* values for individual bilayer leaflets. The theory is based on the concept of elasticity and derives *K_A_* from real-space analysis of local thickness fluctuations sampled in MD simulations. We explore and validate the model on a large set of single and multicomponent bilayers of different lipid composition and sizes, simulated at different temperatures. The calculated bilayer compressibility moduli agree with values estimated previously from experiments and those obtained from a standard computational method based on a series of constrained tension simulations. We further validate our framework in a comparison with an existing polymer brush model (PBM) and confirm the PBM’s predicted linear relationship with proportionality coefficient of 24 using elastic parameters calculated from the simulation trajectories. The robustness of the results that emerge from the new method allows us to revisit the origins of the bilayer mechanical (compressible) thickness and in particular, its dependence on acyl chain unsaturation and the presence of cholesterol.

## INTRODUCTION

Cells exhibit a wide variety of morphologies ranging from discoid or spherical shapes (e.g. erythrocytes and staphylococcus bacteria, respectively), to branched formations with multiple highly curved and flat elongated segments (e.g. nerve axons in the brain, microvilli in the intestines and rod cells in the retina of the eye). A cell’s ability to take on distinct shapes is directly dependent on the flexibility of its bounding plasma membrane (PM), and thus maintaining a certain level of flexibility of the PM is essential to both cell and human physiology. A prominent example of the consequences of deficits in PM flexibility is sickle cell anemia, a disease characterized by a drastic change in the shape and stiffness of red blood cells that leads to their accumulation on vessel walls and blockage of blood flow [1–6].

The PM is composed of two layers, or leaflets, composed of a mixture of various types of amphiphilic lipid molecules, each type with its own set of structural and thermodynamic properties. Despite this complexity, the PM’s flexibility has been successfully modeled quantitatively as a simple elastic sheet with characteristic bending and compressibility constants [7]. While measurements of PM’s elastic properties have been performed directly on live cells, the interpretation of such measurements is still challenging due to the complexity and non-uniformity of the cellular membrane environment [8, 9]. Instead, experiments on less complex, compositionally symmetric model membranes have been utilized to characterize the bilayer’s bending rigidity (*κ_C_*) and area compressibility (*K_A_*) moduli as a function of lipid composition. These mechanical constants quantify, respectively, the energetic cost associated with bending the membrane and stretching/compressing its area, and have thus been used to make successful predictions about biological phenomena, e.g., of the changes in shape of closed lipid vesicles under stress [10–12]. Since the proper function and organization of transmembrane proteins are often regulated by membrane deformations near the protein (e.g. local bilayer bending and thinning or thickening), *K_A_* and *κ_C_* also appear as important parameters in theoretical models quantifying the energetics of protein-membrane interaction [13–15]. All these approaches directly link the elastic properties of membranes to bilayer shape and the sorting of both lipids (e.g., into distinct lipid domains) and proteins (e.g., into local clusters or oligomers) on the surface of a heterogeneous membrane such as the PM.

Various methods exist for measuring *κ_C_* both *in vitro* and *in silico* [16–18], but the equally important compressibility modulus *K_A_* is less well studied. It quantifies the response of membrane area to tension, which under physiological conditions may arise from various perturbations, such as changes in osmolarity across the membrane or the addition of lipids or other molecules to only one of the membrane’s leaflets. Several experimental approaches have been developed to measure *K_A_* in model membranes, and these methods rely on extracting the compressibility modulus from a relationship between systematically varied tension and the resulting bilayer area expansion. Perhaps the most commonly used technique utilizing this approach is the micropipette aspiration of giant unilamellar vesicles (GUVs), which has supplied the largest set of bilayer *K_A_* data available currently [19–21]. The procedure involves imaging a single GUV while applying incremental amounts of suction pressure to it with a micropipette. The tension exerted on the bilayer is calculated directly from the applied pressure while the resulting changes in the bilayer area are inferred from geometrical considerations of the corresponding changes in GUV shape. In an alternative approach, pressure is applied to extruded unilamellar vesicles through osmotic imbalance between the vesicles’ interior and exterior due to solutes such as salt or sugar [22–24]. The ensuing trends in the bilayer structure are monitored from the vesicle diameter measured with techniques such as light scattering or electron microscopy. NMR alone [25] or in combination with X-ray diffraction [26] have also been used to measure *K_A_* of bilayers at low hydration by relating changes in bilayer area to changes in the osmotic pressure of a polymer (e.g. polyethylene glycol) solution. Unfortunately, experimental data on the behavior of many lipids, including lipid mixtures, is still scarce and the availability of the resources needed to make the measurements is often limited. In that respect, a combination of rigorous physics-based simulation and well-calibrated computational tools holds great promise for enabling an otherwise impossible elasticity-based analysis of membrane systems that remain elusive to experimental methods.

With the feasibility of more extensive molecular dynamics (MD) simulations, the area compressibility modulus has been estimated from trajectories of (on average) flat lipid bilayer patches. The classical computational approach is based on the same principle as the experimental methods, i.e., that for small changes in area per lipid (*A_lip_*), tension is linear with direct area expansion. To calculate the bilayer *K_A_*, a series of constrained-area (or tension) simulations is performed and the value emerges from the slope of the best-fit line through the data of ln (*A_lip_*) vs surface tension. While the estimated moduli are typically in good agreement with experimental estimates, the analysis of one lipid composition requires multiple simulations, which makes this approach very expensive computationally. An alternative computational strategy that circumvents this requirement uses the equilibrium thermal fluctuations of the bilayer at constant zero tension instead. In this spirit, *K_A_* is estimated from a single simulation trajectory utilizing the equilibrium expression 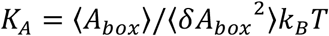, where *A_box_* represents the lateral area of the simulation box, *k*_8_ is the Boltzmann constant, *T* is temperature and 〈 · 〉 denotes ensemble average (see Methods section below). Since the analysis is directly related to the fluctuations in the simulation box, the modulus exhibits a strong dependency on the thermodynamic phase behavior of the bilayer (which is directly related to the relaxation rate of its lateral area), system size, and the corresponding length of the simulation trajectory. Importantly, no existing computational or experimental methods allow for calculation of *K_A_* of individual bilayer leaflets, which is a prerequisite for the quantification of the energy of local leaflet distortions in an asymmetric bilayer.

Here, we report a novel computational methodology that overcomes the aforementioned shortcomings in the calculation of area compressibility, and obtains the *K_A_* moduli of a bilayer and of each of its leaflets from a single MD trajectory. Like the existing methods, we take advantage of thermal fluctuations, but express the compressibility modulus as a function of leaflet thickness instead of bilayer area. Following our recent success in calculating each leaflet’s bending rigidity from real-space analysis of local splay fluctuations [18], we base our method for *K_A_* estimation on sampling the leaflet thickness locally, and estimate the corresponding probability distribution and potential of mean force (PMF) profile as a function of changes in thickness. Finally, the *K_A_* is extracted from a quadratic fit of a small region of the PMF around the global minimum, according to the elastic energy of stretching (see Methods). We show that for a large set of single and multicomponent bilayers, the compressibility moduli we calculate with the new method are in excellent agreement with the ones reported from experiments *in vitro*, or calculated with alternative computational approaches. We find that the *K_A_* values obtained with our framework are less sensitive to bilayer size and simulation length due to the local nature of the analysis. We further validate our approach by reproducing the linear relationship between bilayer thickness, *K_A_* and *κ_C_* in the polymer brush model (PBM) [20], using mechanical constants calculated from the simulation trajectories. This analysis lets us revisit the definition of the bilayer mechanical thickness and clarify observed discrepancies reported in the literature with respect to PBM’s predictions [17, 27, 28].

## METHODS

Here we present the theoretical framework and details of the new method for the calculation of the area compressibility moduli of a bilayer and each of its leaflets, based on the analysis of trajectories from molecular dynamics simulations. Section I contains an overview of compressibility calculation from area fluctuations, followed by the formulation for the calculation of individual local leaflet moduli. In Section II we re-express the theory in terms of local leaflet thicknesses and provide a detailed methodological description of the new computational framework in order to address some of the challenges presented by using area in the formulation developed in the previous section. Section III presents details of the simulation trajectories used for the application of the method, as well as a summary of the corresponding analysis.

### I. Compressibility modulus from area fluctuations

#### Bilayer compressibility

Following Helfrich’s formalism [29], we treat the bilayer as a two-dimensional elastic sheet with mechanical constants describing its modes of deformation. For small equilibrium fluctuations around the free energy minimum, each deformation mode is associated with an elastic energy that is approximated by a quadratic function of the relevant deformation, analogous to Hooke’s law. For changes in area, the elastic energy *E* of stretching/compressing a bilayer patch with equilibrium area *a*_0_is given as (see Eq. 1 in [30]):

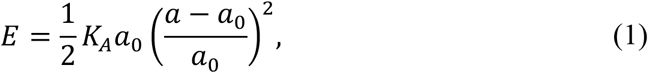

where *a* is the deformed area and *K_A_* is the bilayer area compressibility modulus. Assuming that bilayer area stretching/compression are independent degrees of freedom in the context of the full energy functional describing bilayer mechanics (cf. bending or tilt), *K_A_* can be obtained from the bilayer’s thermally excited area fluctuations [31]. Specifically, from the equipartition theorem, 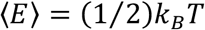 where *k*_8_ is Boltzmann constant and *T* is temperature, it follows that:

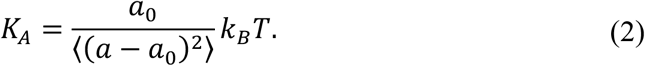

This is the expression commonly used to obtain *K_A_* from MD simulations of a bilayer by sampling fluctuations in the lateral area of the simulation box *A_box_* [17, 31, 32]. Since *E* is the energy of a deformed state described by Δ*a*/*a*_0_ = (*a* – *a*_0_)/*a*_0_, from statistical mechanics it also follows that the probability of this state can be written as:

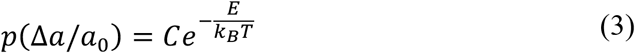

where *C* is a constant. Rearrangement of Eq. 3 leads to an alternative equality from which *K_A_* can be calculated provided the probability distribution of the deformed states is known:

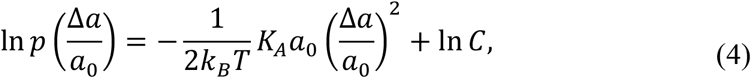

which can be written as:

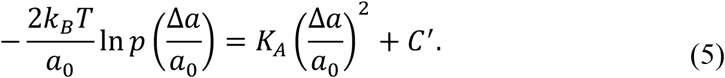

While both Eqs. 2 and 5 are equivalent upon sufficient sampling of area fluctuations, Eq. 2 in which *K_A_* is inversely proportional to the mean square area fluctuations, is more sensitive to outliers and deviations from the elastic regime (see Eqs. 12–13 and discussion afterwards) when used for *K_A_* estimation. In contrast, Eq. 5 relies on the distribution of deformations around the mean and can thus provide a more robust approximation of the area compressibility modulus.

#### Leaflet compressibility

The area compressibility modulus of a bilayer quantifies the total energy of membrane deformation and can be used to infer the energetics of deforming individual bilayer leaflets in *symmetric* bilayers whose two leaflets are assumed to behave in the same way. However, in an *asymmetric* membrane, the two leaflets can have very different lipid compositions with potentially different energetic costs for stretching/compression that cannot be simply inferred from the elastic properties of the bilayer. To enable the analysis of these more general (and physiologically relevant) systems, we sought a formulation that would yield the area compressibility modulus of each leaflet of the membrane independently. Specifically, the goal was to obtain leaflet compressibility from area fluctuations in the spirit of the above-described theory (Eqs. 2–5).

Globally (e.g. considering the entire simulated bilayer patch), the area fluctuations of the two leaflets are identical and equal to the area fluctuations of the whole bilayer. Therefore, the apparent leaflet compressibility moduli calculated from area fluctuations at that scale would always appear the same, masking any potential differences in the inherent mechanical properties of the leaflets. In order to extract these differences and find the effective local leaflet moduli, we perform the analysis on a length scale much smaller than the global bilayer area. In particular, we view each leaflet as a collection of more than one parallel elastic blocks that are made of the same material (i.e. have the same compressibility modulus). Within a leaflet, it is assumed that all blocks have the same average area (e.g. the average area of a lipid) but can have different instantaneous areas and their area fluctuations are weakly coupled. Due to its elastic nature, the deformation energy of a block has the same form as Eq. 1 and its compressibility modulus (which is the effective local leaflet modulus) can be obtained accordingly from its area fluctuations through Eq. 2 or Eq. 5.

In order to relate the effective local leaflet moduli to the bilayer’s compressibility, we express the energy of bilayer stretching/compression as a function of the stretching/compression of the individual leaflet blocks. If we denote the instantaneous global areas of the two leaflets with *A_x_* and *A_y_* and the instantaneous and average local areas of their elastic blocks with *a_xi_*, *a_yi_* and *a_xo_*, *a_yo_* respectively (*i* being the identity of the block), the second order approximation of the bilayer energy can be written as:

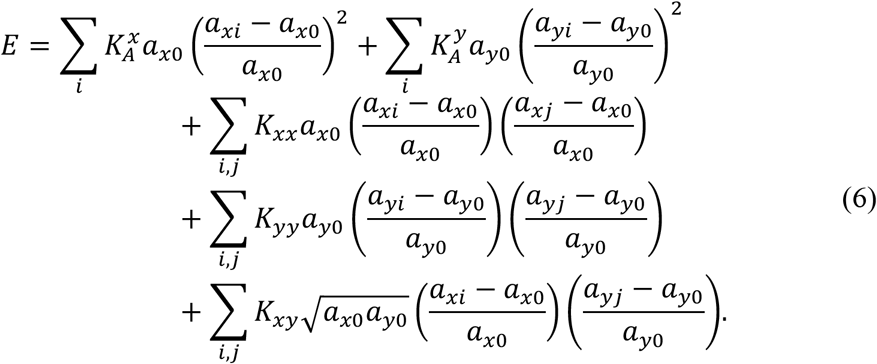

The first two terms in Eq. 6 represent summations over the deformation energies of the individual elastic blocks in the two leaflets, the next two terms are the corresponding inter-block correlations within each leaflet, and the last term quantifies the correlations between blocks from different leaflets. Each term has its characteristic modulus, and 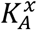 and 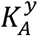 in particular are the effective local leaflet compressibility moduli.

Since the bilayer area *A* is *A* = *A_x_* = *A_y_* = (*A_x_* + *A_y_*)/2, *A_x_* = ∑*_i_a_xi_* and *A_y_* = ∑*_i_a_yi_*, we can express the variance of *A* as a sum over variances and covariances of the local areas:

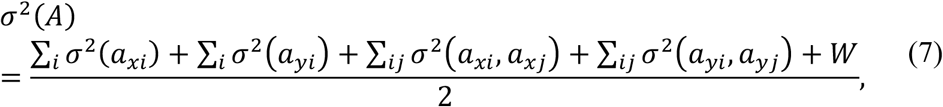

where *W* = ∑_*ij*_σ^2^(*a_xi_*, *a_yj_*). Since the global areas of the two leaflets are constrained to be the same, their variances are the same and consequently, the sum of the interleaflet correlations is 0, i.e. *W* = 0.

In addition, 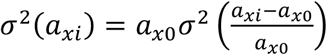 and at the same time 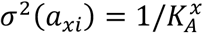 (as in the second order approximation of the energy the multivariate Boltzmann distribution becomes the multivariate normal distribution). If *n* denotes the number of blocks, Eq. 7 then simplifies to:

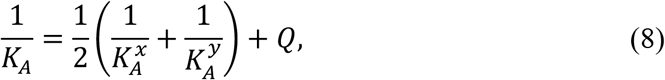

where *Q* is the average sum of the inter-block correlations within each leaflet:

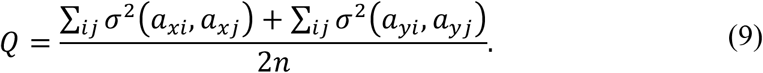

While these inter-block correlations within a leaflet can generally deviate from 0, when the local areas (elastic blocks) are small the correlations can be both positive and negative (representing the fact that the blocks can undergo stretching/compression deformations in two dimensions) and we find that they cancel each other out in the respective sums (see section S.1 in SM). As a result, we assume *Q* ≈ 0 and arrive at the final relationship between the bilayer *K_A_* and the local compressibility moduli, 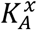 and 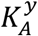, of the two leaflets:

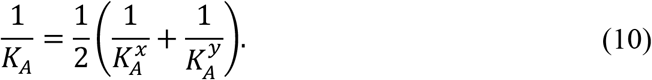

Note that the derivation above assumes that the two leaflets have the same number of elastic blocks, however Eq. 10 holds even in the general case when these numbers are different (as for membranes with asymmetric lipid composition; see section S.1 in SM for the extended derivation).

It is important to note that in our formulation a local leaflet *K_A_* in a symmetric bilayer has the same magnitude as the bilayer *K_A_* and therefore should be treated differently from the *global* leaflet compressibility moduli often referred to in the literature as 1/2 the bilayer *K_A_* [33]. The latter are based on a model in which the global area changes in the two leaflets are the same due to the constraints on the bilayer geometry, but are uncoupled, and thus the elastic energy (and consequently, the *K_A_*) of deforming each leaflet is half the energy (*K_A_*) of deforming the bilayer. In contrast, 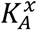 and 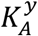 capture the *local* properties of the leaflets, which are affected by the global constraint on area only indirectly and thus reveal features that are more specific of the leaflets themselves. From Eqs. 6–10 it further follows that the harmonic mean of the local leaflet moduli gives the bilayer *K_A_*, which as we show in Results quantitatively matches experimentally measured bilayer moduli for various membrane systems.

### II. Compressibility modulus from thickness fluctuations

The theoretical formulation presented in the previous section allows the calculation of an effective local leaflet compressibility modulus from area fluctuations. To capture the individual leaflet properties when calculating 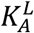 (where *L* can be *x* or *y*), given the outlined considerations (e.g. the cancelation of the *Q* term in Eq. 8), we chose to analyze the fluctuations of the smallest local unit area that is characteristic for leaflet *L*, that is the average area per lipid in the leaflet. In this way, the two leaflets in a symmetric bilayer will have the same local unit area, as expected, while in an asymmetric bilayer they may be different. We thus seek a local description of instantaneous leaflet area that would allow for ample sampling of area fluctuations, while treating each leaflet independently from the other leaflet.

Since the definition and calculation of local leaflet areas is rather challenging [31, 34], we assume volume conservation to relate deformations in local area to deformations in local thickness, and then estimate the coefficients from thickness fluctuations. Specifically, let *a^L^* and *t^L^* be the instantaneous local area and thickness of a leaflet, and 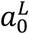 and 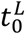 are their corresponding equilibrium values. Assuming that 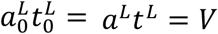 where *V* is a constant, we can express the energy of stretching/compressing leaflet *L*, *E^L^*, as a function of *characteristic* changes in thickness instead of *relative* changes in area:

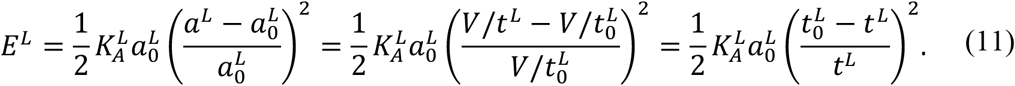

Consequently, Eqns. 2 and 5 become:

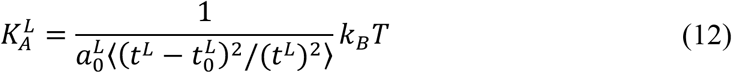

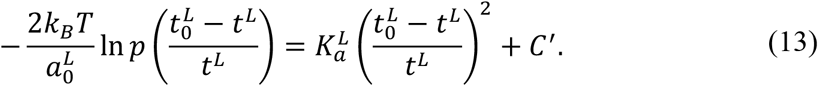

While both Eqns. 12 and 13 can be used to obtain 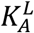 in theory, practical (numerical) considerations render Eq. 13 more suitable (see section S.2 in SM for comparison between the two approaches). Our formal framework is therefore centered on extracting 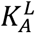 from Eq. 13 through the following steps:

1. Calculate local leaflet thicknesses at different chain depths within the bilayer.
2. From all possible definitions of leaflet thickness, identify the one that is suitable for the calculation of 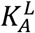.
3. Use Eq. 13 to obtain 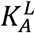.

In the following, we present details of the above three stages in our algorithm.

#### 1. Calculating local leaflet thicknesses from simulations

Over the course of a simulation trajectory, the thickness of each bilayer leaflet is laterally inhomogeneous and fluctuates around its equilibrium value 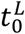 as a result of thermal motions. To construct the probability distribution of thickness changes in Eq. 13, *p*(Δ*t*/*t^L^*), by sampling local fluctuations, we take a grid-based approach and calculate the leaflet thickness at different grid points on the leaflet surface. In a continuum representation, a leaflet can be viewed as a stack of layers with each layer being a surface made of a particular lipid atom (*ς*), e.g. a phosphate surface (ς = *P*), a first glycerol carbon surface (*ς* = *C1* using CHARMM36 atom naming scheme), a first sn-1 carbon surface (*ς* = *C21*), a first sn-2 carbon surface (*ς* = *C31*) and so on (see Fig. S1). To calculate the leaflet thickness at a grid point (*x,y*) we first find the height of each of these surfaces at this grid point by performing interpolation on the corresponding atomic z-coordinates as follows:

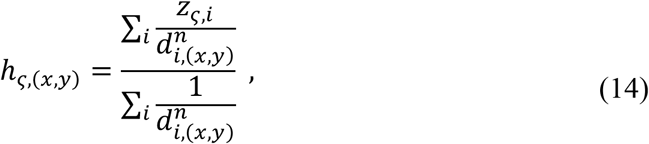

where *h_ς,_*_(*x,y*)_ is the height of the *ς*-surface (i.e. the surface made of atom type *ς* at grid point (*x*,*y*))), *z_ς,i_* is the z-coordinate of atom *i*, *d_i_*_,(*x*,*y*)_ is the 2D distance between atom *i* and (*x*,*y*), *n* is interpolation order, and the summations are done over all leaflet atoms *i* of type *ς* (note that the atoms on individual lipid chains have unique atom names, thus each lipid has at most one atom of type *ς*). Since a lipid can have multiple chains and the heights in Eq. 14 are calculated separately for each carbon atom on each chain, we simplify the analysis by averaging the corresponding surface heights across all lipid chains:

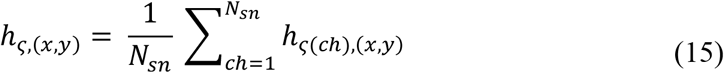

where *h_ς_*_(*Ch*), (*x,y*)_ is the height of the surface calculated from the *ς* carbon on the *Ch* chain, and *N_sn_* is the number of lipid chains. Eq. 15 can also be extended to lipids whose chains have different lengths (see Section S.3 in SM). Thus, a single ‘average chain’ is created at each grid point and used in the subsequent analysis. While this approach works well for most bilayers, we found that for lipids like SAPE (see Simulation details below for lipid name abbreviations) which has one fully saturated and one highly-unsaturated chain, this averaging can become problematic for the subsequent analysis that is based on correlations of the resulting heights (described in the next section). Empirically, we found that the problem is alleviated when prior to the averaging in Eq. 15, each double bond and its preceding carbon are represented by a single data point with an instantaneous height equal to the average interpolated height across the 3 carbons. Thus, each double bond effectively reduces the unsaturated chain length (or the number of surfaces defining the unsaturated chain) by 2 carbons. As this procedure is more general and at the same time does not affect the results for the bilayers for which Eq. 15 can be applied, it was integrated in the methodological framework.

The leaflet thickness at the level of the *ς*-surface, denoted τ_*ς*_, is defined as the difference between the height of the *ς*-surface and the height of the lowest-situated surface at the grid point (usually, the surface of the terminal methyl carbons):

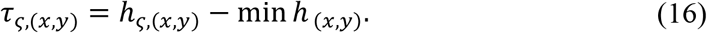

The interpolation order *n* in Eq. 14 determines the contribution to *h_ς_*_(*x,y*)_ of the atoms closest to (*x,y*) relative to those that are further apart, i.e., the higher the *n*, the larger the effect from nearby atoms, and the lower the *n*, the larger the effect from all atoms. Hence, *n* is related to the effective number of atoms (lipids) that are being averaged, and consequently to the equilibrium area 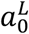 in Eq. 13 (i.e. the area whose thickness fluctuations are effectively being analyzed).

Since atoms are weighted differently in the interpolation, we estimate 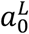 by first using the denominator in Eq. 14 to approximate the *effective* number of lipids being averaged, and then multiplying this number by the average area per lipid in the leaflet, 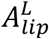. When *n* = 2 the denominator in Eq. 14 is approximately 1, conveniently setting 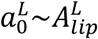, and we therefore use *n* = 2 and 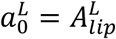 in the subsequent analysis (see Section S.4 in the SM for the derivation, and Fig. S2 for a demonstration of the invariability of the results with *n*). As described in the SM (Section S.5), the interpolated thicknesses (Eq. 16) preserve the product of local area and thickness (i.e. the assumption underlying Eq. 11), which further establishes their suitability for the calculation of 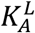.

#### 2. Identifying the relevant thickness for fluctuations analysis

The first step of the method described above allows us to calculate local thicknesses at different surfaces (i.e. different depths) within the leaflet. Naturally, surfaces at heights near the water/hydrocarbon interface fluctuate less due to interfacial tension, while those further down into the leaflet fluctuate more, due to the increased fluidity of the lipid chains around the bilayer midplane. The height (and consequently thickness) fluctuations in a leaflet therefore fall roughly into two categories: ones that are relatively suppressed, and ones that are dominated by relatively unconstrained motion of the lipid chains. The former would tend to overestimate 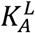 while the latter would tend to underestimate it. The fluctuations of the surface lying at the intersection of these two regimes will thus be the most suitable from the elasticity considerations to obtain a reliable estimate of 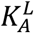. We term the thickness at the level of this surface the *relevant thickness for* 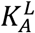 and proceed to identify its location within a bilayer leaflet.

The location of the atomic surface corresponding to the relevant thickness may differ in different membranes due to various degrees of bilayer fluidity. We have therefore developed a general algorithm for identifying the surface relevant for 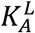 calculations for an arbitrary lipid membrane. Specifically, we examine the correlation between the height fluctuations of a particular surface ς and those of a reference surface (*RS*) close to the water-hydrocarbon interface (in our case, the *RS* is the surface of the first acyl chain carbon atom that is not connected to oxygen). Fig. 1 shows a typical behavior of this correlation, *r*(*ς*) = *r*(*h_RS_*, *h_ς_*), (here, for the top leaflet of a DPPC bilayer) as the distance, *d*(*ς*), between a surface ς and the reference surface increases 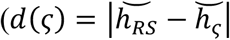; where 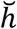) denotes median of the distribution of local heights). At small *d*(*ς*), the correlation drops slowly and linearly with distance (Fig. 1, red solid line), implying a regime of suppressed thickness fluctuations (i.e. fluctuations of the atoms in this segment of the chain are strongly coupled to one another). At larger *d*(*ς*) distances, the *r*(*ς*) vs. *d*(*ς*) profile strongly deviates from the initial linear behavior as *d*(*ς*) increases more slowly while *r*(*ς*) decreases more rapidly, characteristic of the more fluid region of the bilayer in which the lipid chains exhibit greater flexibility and intercalate with the chains of the opposing leaflet. Given the two well-defined regimes, we choose the first point outside of the linear regime (Fig. 1, yellow dashed line) to represent the leaflet whose thickness fluctuations can be used to extract 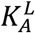. We identify this surface *ς*_s_ following the algorithm outlined on the right hand side box in Fig. 1. For DPPC the surface *ς*_s_ is at the 10th carbon (shown in yellow in Figs. 1 and S1). Interestingly, for this and all other bilayers that we examined, *ς*_s_ appears to be located right around the region within the leaflet where the density of the opposing leaflet vanishes (i.e. just outside the interleaflet interdigitation zone, see Fig. S3).

**Figure 1.**
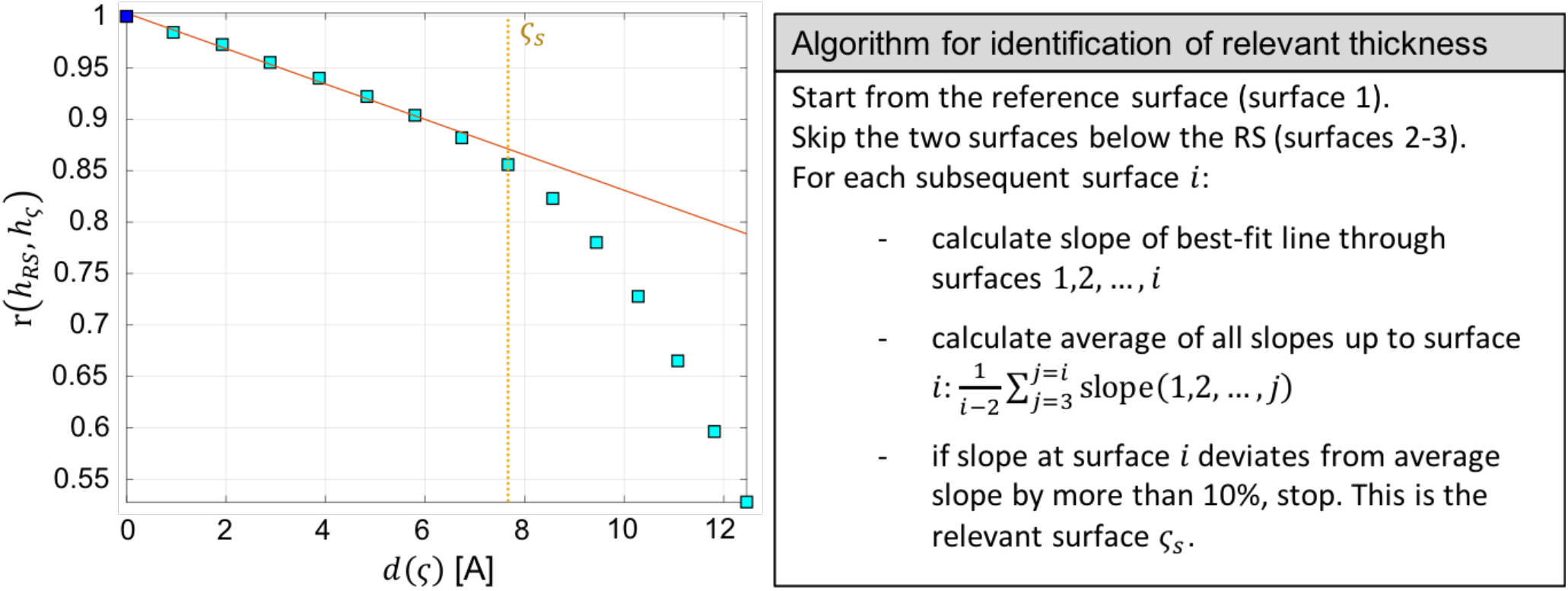
Identification of relevant thickness for calculation of 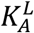. Left, the correlation of the height fluctuations of a surface *ς* with the height fluctuations of a *reference surface* RS (first acyl chain carbon atom not attached to oxygen, shown in blue here and in Fig. S1) is plotted as a function of distance from RS for the top leaflet of a DPPC membrane. Right, outline of the algorithm used to identify the relevant surface *ς_s_* from the data (denoted by a yellow dotted line on the plot). The solid red line shows the corresponding best-fit line through all points preceding *ς_s_*. A sample representation of the relevant surface can be seen in yellow in Fig. S1.

#### 3. Calculating 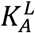from local thickness fluctuations

Having identified *ς_s_*, we calculate the local thickness 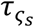 at every grid point in every frame of the trajectory and estimate its probability distribution using kernel density estimation (Fig. 2A). From this distribution, we estimate *t*_0_ (defined here and throughout as the most probable thickness, i.e. at the peak of the distribution), and the left hand side of Eq. 13, i.e. the Potential of Mean Force (PMF), as shown on Fig. 2B. The characteristic asymmetric shape of the PMF is consistent with the free energy vs. area per molecule profile predicted by Ben-Shaul from theoretical considerations of lipid chain packing [35] and arises from the relative ease of deforming the membrane upon thickness contraction (area expansion, increase in entropy) compared to thickness expansion (area contraction, decrease in entropy). To find the leaflet compressibility, we identify a small region around *t*_0_ (between 5% and 7% of *t*_0_) where the PMF is closest to a normal distribution. We then fit a quadratic function to the PMF in this region and obtain 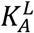 from the quadratic coefficient in the best fit (see Section S.6 in SM for a step-by-step outline of the procedure). Errors are calculated with a two-dimensional bootstrapping approach over both time and space as described in Section S.7 in SM.

**Figure 2.**
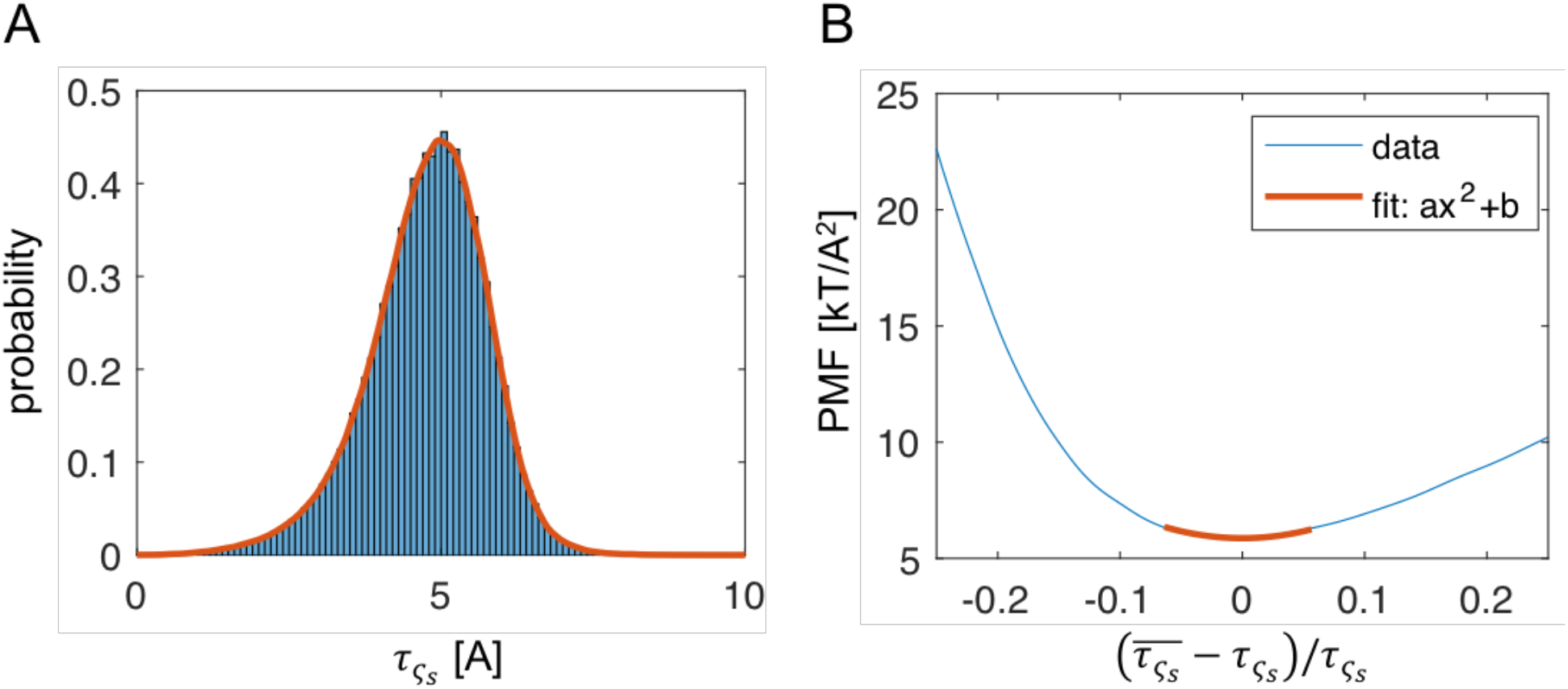
Calculation of 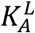 from local thickness fluctuations. A, Probability distribution of the relevant thickness, 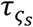, constructed from the time evolution of the local interpolated thicknesses for the top leaflet of a DPPC bilayer (blue). The distribution is smoothed for subsequent analysis by using a kernel density (red). B, The left-hand side of Eq. 16 is plotted as a function of characteristic changes in the local thickness. The PMF in the region of thicknesses within 6% of the mean thickness is fit to a function of the form *y* ~ *ax*^2^ + *b* (see text for details). 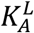 is obtained from the quadratic coefficient *a* = 238 mN/m of the best fit (shown in red).

### III. Simulations and analysis

#### Simulation details

Table S1 lists information for all simulated bilayers. The following lipid names are abbreviated as shown in the parenthesis: 1,2-dilauroyl-*sn*-glycero-3-phosphocholine (DLPC, di12:0), 1,2-dimyristoyl-*sn*-glycero-3-phosphocholine (DMPC, di14:0), 1,2-palmitoyl-*sn*-glycero-3-phosphocholine (DPPC, di16:0), 1,2-dilinoleoyl-*sn*-glycero-3-phosphocholine (DLiPC, di18:2), 1,2-dioleoyl-*sn*-glycero-3-phosphocholine (DOPC, di18:1(cis)), 1-stearoyl-2-oleoyl-*sn*-glycero-3-phosphocholine (SOPC, 18:0,18:1), 1,2-dielaidoyl-*sn*-glycero-3-phosphocholine (DEPC, di18:1(trans)), 1-palmitoyl-2-oleoyl-*sn*-glycero-3-phosphocholine (POPC, 16:0,18:1), 1-palmitoyl-2-oleoyl-*sn*-glycero-3-phosphoethanolamine (POPE, 16:0,18:1), 1-stearoyl-2-diarachidonoyl-*sn*-glycero-3-phosphoethanolamine (SAPE, 18:0,20:4), 1,2-dioleoyl-*sn*-glycero-3-phospho-(1’-rac-glycerol) (DOPG, di18:1(cis)), 1’,3’-bis[1,2-dioleoyl-*sn*-glycero-3-phospho]-*sn*-glycerol (TOCL, tetra18:1), N-palmitoyl-D-*erythro*-sphingosylphosphorylcholine (PSM), cholesterol (Chol). As indicated in Table S1, some of the bilayers were taken from Ref. [18]. All remaining bilayers were constructed with CHARMM-GUI [36–38] and simulated with NAMD [39] using the CHARMM36 force field [40, 41]. Initial equilibration was carried out with CHARMM-GUI’s protocols. Following equilibration, the simulations were performed with a 10-12 Å cutoff for non-bonded and short-range electrostatic interactions, and Particle Mesh Ewald with grid spacing of 1 Å for long-range electrostatics. Vdw force switching was turned on. Temperature was controlled with a Langevin thermostat while constant pressure was maintained with NAMD’s Langevin piston Nose-Hoover method with a 200 fs period and 50 fs decay. All simulations were run with a time step of 2 fs, *rigidbonds* set to all, and both atomic coordinates and velocities were output every 20 ps.

#### Simulation analysis and method implementation

All bilayer properties were estimated from the last segments of the trajectories over which the lateral area of the simulation box was considered converged, as determined with a method based on maximizing the number of effectively uncorrelated data points [42]. Table S1 lists the total simulation time for each system and the length of the trajectory segments used for the analysis. All trajectories were centered prior to analysis, such that the mean z position of the terminal methyl carbons on all lipids was set to 0.

The interpolated heights, as described in Step 1 of the calculation of leaflet compressibility from thickness fluctuations, were calculated for each leaflet with a modified version of VMD’s MEMBPLUGIN [43] and sampled on an 8×8 Å^2^ square grid. All subsequent analysis (outlined step-by-step in Section S.6 in SM) was performed with MATLAB. Number density profiles were calculated with the Density Profile tool in VMD [44] and acyl chain order parameter profiles were obtained with in-house Tcl and MATLAB scripts. All code for calculation of the area compressibility moduli is available upon request.

#### Lateral pressure profile calculation

Lateral pressure profiles were calculated from the last ~100 ns of the centered simulation trajectories. The calculation was done with NAMD using stored instantaneous atomic coordinates and velocities. Each system was divided into slabs of approximately 0.8 Å thickness and lateral area equal to the area of the simulation box (all slabs in a given frame of the trajectory have equal volume). The total lateral pressure in each slab was the sum of the independently obtained Ewald and non-Ewald pressure contributions. The x, y and z dimensions of the grid size used for calculating the Ewald contribution were all equal and less than half the z dimension of the simulation box (typically, 30 Å). Due to the known limitations of the Harasima algorithm with PME electrostatics implemented in NAMD for the discretized pressure calculation, the normal component of the pressure tensor in each slab, *p*_v_, was set to 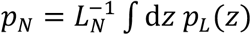 where *L_N_* is the length of the simulation box in the direction of the bilayer normal, and *p_L_*(*z*) is the lateral (or tangential) pressure in slab z [45, 46]. The total pressure in slab z was then calculated as *p*(*z*) = *p_L_*(*z*) − *p_N_*.

#### Treatment of Chol-containing membranes

At relatively small mole fractions (up to 0.3-0.35) of cholesterol (Chol) in the bilayers, the expected effect of Chol on the bilayer structural properties (average area and volume per lipid) is mainly a *condensing* effect on the non-Chol lipids [34, 47]. It is reasonable, therefore, to assume that Chol’s effect on the area compressibility modulus at such small Chol mole fractions will be indirect and captured in the *K_A_* calculated from the non-Chol components only, as detailed in Section S.3 in the SM. However, as shown (see, for example, Ref. [47]), at higher mole fractions (above 0.35) the distribution of Chol’s tilt angles becomes narrower and moves closer to zero, indicating that its motion is more restricted and the molecule is more parallel to the bilayer normal. In this regime Chol is likely to contribute directly to the leaflet compressibility (i.e. compression of the bilayer would involve compression of the Chol molecules themselves), and its effect needs to be considered explicitly. This is achieved by assuming that the ratio of the area compressibility calculated by considering only the non-Chol components 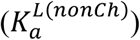 and all components 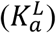 is the same as the fraction (*s_f_*) of the surface area occupied by the non-Chol lipids, i.e. 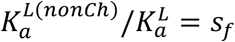 (where 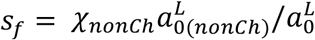. From here 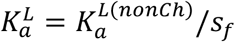. This correction to 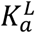 is used only for bilayers with Chol mole fractions > 0.35, and is found to produce a gradual increase in the bilayer compressibility modulus, consistent with experimental measurements (see Fig. 3 and discussion below).

**Figure 3.**
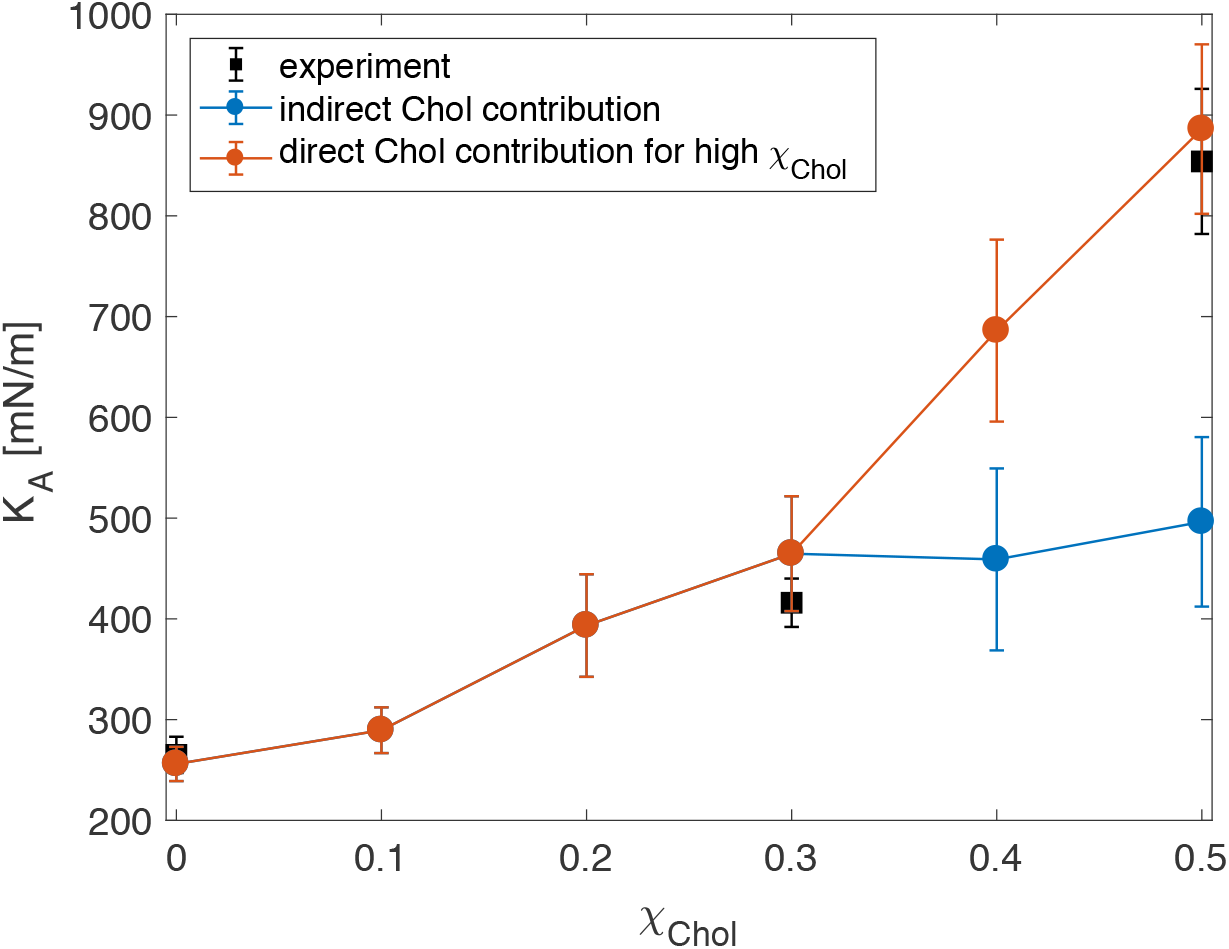
Bilayer area compressibility for DOPC/Chol mixtures with increasing amount of cholesterol. The bilayer *K_A_* was calculated either by considering only the non-Chol lipids for all Chol mole fractions *χ_Chol_* (blue symbols) or the non-Chol lipids for *χ_Chol_* ≤ 0.3 and the direct contribution of Chol to the bilayer compressibility for *χ_Chol_* > 0.3 (red symbols, see Simulations analysis and method implementation section for details). Experimentally determined compressibility moduli are shown in black. The red and blue lines are guides to the eye to the respective data points.

## RESULTS AND DISCUSSION

### Validation of the new method for quantifying compressibility moduli from local thickness fluctuations (LTF)

The canonical method for calculating bilayer area compressibility from MD simulations involves performing a series of NPγT simulations (e.g., [31]). In each simulation a non-zero tension γ is applied in the x-y plane and the bilayer *K_A_* is obtained by analyzing the linear relationship between the applied tension and the resulting area expansion [31]. Since this methodology directly mimics the micropipette aspiration technique [48], it is expected to reproduce closely the experimental results provided that the properties of the simulated bilayers are correctly captured by the underlying force field. Table 1 shows the calculated moduli from NPγT simulations of several single-component lipid membranes and the corresponding experimental measurements (see also Table S2). There is indeed a very good agreement between the two, verifying that the simulations are well-converged and able to reproduce the known values of *K_A_* using standard techniques. Therefore, these and other single and multicomponent symmetric bilayers served as controls for the validation of our new LTF method.

**Table 1.** Area compressibility of compositionally symmetric single component bilayers calculated with different methods: local thickness fluctuations (LTF), constrained tension (NPγT) simulations and box (i.e. bilayer) area fluctuations. Where available, experimentally measured moduli are shown (Exp) with references to the respective literature sources. All moduli are given in units of mN/m. The LTF bilayer *K_A_* was calculated with Eq. 10. LTF errors for the leaflets were calculated with a 2-dimensional bootstrap method as described in Section S.7 in SM and propagated to obtain the error on the bilayer *K_A_*. The 95% confidence interval for the values obtained from a linear fit of tension versus area expansion in NPγT simulations (Table S2) are shown in brackets. For lipid name abbreviations see Simulation details section in the text. DOPC rep 1 and DOPC rep 2 represent two independently constructed and simulated replica trajectories of the DOPC bilayer. The four POPC bilayers labeled 1 to 4 differ in their size, simulation temperature and salt concentration, as shown in Table S1.

| Bilayer | LTF | | | Exp | NPγT simulations | Box Area Fluctuations |
| --- | --- | --- | --- | --- | --- | --- |
|  | top | bottom | bilayer |  |  |  |
| DLPC | 258 ± 28 | 214 ± 22 | 234 ± 17 | | | 213 ± 20 |
| DMPC | 236 ± 20 | 236 ± 22 | 236 ± 15 | 234<sup>a</sup> ± 23 | 235<br>[172, 297] | 263 ± 23 |
| DPPC | 238 ± 34 | 274 ± 34 | 255 ± 24 | 231<sup>b</sup> ± 20 | 223<br>[188, 257] | 183 ± 20 |
| DLiPC | 200 ± 26 | 202 ± 22 | 201 ± 17 | 247<sup>a</sup> ± 21 | 237<br>[189, 285] | 261 ± 23 |
| DOPC [full] | 240 ± 20 | 274 ± 28 | 256 ± 17 | 265<sup>a</sup> ± 18<br>310<sup>c</sup> ± 20 | 253<br>[211, 295] | 246 ± 20 |
| DOPC [part] | 266 ± 32 | 260 ± 26 | 263 ± 21 | | | 313 ± 33 |
| DOPC rep 1 | 246 ± 24 | 256 ± 26 | 251 ± 18 | | | 223 ± 31 |
| DOPC rep 2 | 272 ± 28 | 272 ± 68 | 272 ± 37 | | | 274 ± 26 |
| SOPC | 260 ± 24 | 232 ± 32 | 245 ± 21 | 235<sup>a</sup> ± 14<br>290<sup>c</sup> ± 6 | | 236 ± 21 |
| DEPC [full] | 212 ± 24 | 214 ± 22 | 213 ± 16 | 229<sup>a</sup> ± 12 | | 204 ± 18 |
| DEPC [part] | 240 ± 38 | 186 ± 42 | 210 ± 30 | | | 321 ± 53 |
| POPC 1 | 206 ± 32 | 168 ± 20 | 185 ± 18 | 255<sup>d</sup> ± 75 | 214<br>[134, 293] | 220 ± 17 |
| POPC 2 | 236 ± 26 | 186 ± 28 | 208 ± 20 | | | 172 ± 30 |
| POPC 3 | 258 ± 28 | 234 ± 32 | 245 ± 22 | | | 250 ± 23 |
| POPC 4 | 214 ± 38 | 198 ± 36 | 206 ± 26 | | | 236 ± 22 |
| POPE | 242 ± 28 | 218 ± 15 | 229 ± 30 | 233<sup>e</sup> | 285<br>[260, 310] | 291 ± 26 |
| SAPE | 230 ± 12 | 208 ± 20 | 218 ± 25 | | | 265 ± 14 |
| DOPG | 236 ± 13 | 230 ± 14 | 233 ± 19 | | | 220 ± 26 |
| TOCL | 224 ± 8 | 238 ± 7 | 231 ± 11 | | | 254 ± 20 |
| PSM | 344 ± 20 | 286 ± 20 | 312 ± 29 | | 324<br>[251, 396] | 499 ± 42 |

<sup>a</sup> Ref. [20], <sup>b</sup> Ref. [61], <sup>c</sup> Ref. [21], <sup>d</sup> Ref. [49], <sup>e</sup> Ref. [62]

Tables 1 and 2 list the compressibility moduli for each leaflet and for the bilayer obtained from LTF analysis of the equilibrated trajectories (see Table S1 for a detailed description of all simulated bilayers). The bilayer *K_A_*’s obtained from the LTF method are in excellent agreement with those measured experimentally or obtained from NPγT simulations (Table 1), indicating that our new approach reproduces the accuracy of existing methods when applied to symmetric bilayers. We note, however, that while the confidence intervals associated with the linear fits from the NPγT simulations are large, the LTF errors calculated with a 2D bootstrapping algorithm (as described in Section S.7 in SM) are much smaller and comparable to the experimental uncertainties. (The experimental uncertainty for POPC is notably larger than the rest because the measurement was obtained with a different infrared (IR) linear dichroism-based method [49].)

**Table 2.** Area compressibility of compositionally symmetric multicomponent bilayers calculated with the three different methods described in the caption of Table 1. For lipid name abbreviations see Simulation details section in the text.

| Bilayer | LTF method |  |  | Exp | Box Area<br>Fluctuations |
| --- | --- | --- | --- | --- | --- |
|  | top | bottom | bilayer |  |  |
| POPC/Chol<br>70/30 | 862 ± 92 | 676 ± 126 | 757 ± 87 | 673<sup>a</sup> | 562 ± 87 |
| DOPC/Chol<br>90/10 | 276 ± 36 | 306 ± 24 | 290 ± 23 |  | 260 ± 27 |
| DOPC/Chol<br>80/20 | 466 ± 62 | 340 ± 68 | 393 ± 51 |  | 338 ± 34 |
| DOPC/Chol<br>70/30 | 544 ± 74 | 406 ± 76 | 465 ± 57 | 416<sup>a</sup> ± 24 | 532 ± 75 |
| DOPC/Chol<br>60/40 | 804 ± 118 | 598 ± 120 | 686 ± 90 |  | 829 ± 76 |
| DOPC/Chol<br>50/50 | 956 ± 140 | 826 ± 102 | 886 ± 84 | 854<sup>a</sup> ± 72 | 1011 ± 99 |
| DPPC/Chol<br>80/20 | 2916 ± 364 | 3368 ± 436 | 3126 ± 281 |  | 1968 ± 208 |
| POPE/POPG<br>70/30 | 194 ± 22 | 228 ± 28 | 210 ± 17 |  | 193 ± 19 |
| POPC/POPS<br>70/30 | 350 ± 40 | 318 ± 34 | 333 ± 26 |  | 360 ± 43 |
| POPE/POPS<br>70/30 | 366 ± 40 | 332 ± 50 | 348 ± 33 |  | 322 ± 69 |
| DMPC/POPC<br>10/90 | 238 ± 22 | 188 ± 32 | 210 ± 22 |  | 258 ± 19 |
| DMPC/POPC<br>43/57 | 264 ± 26 | 230 ± 32 | 246 ± 21 |  | 224 ± 19 |
| DMPC/POPC<br>75/25 | 282 ± 24 | 244 ± 38 | 262 ± 24 |  | 250 ± 19 |

<sup>a</sup> Ref [19]

Table 1 also lists the corresponding compressibility moduli calculated from the same equilibrated portions of the trajectories, but from lateral bilayer area fluctuations using Eq. 2. While these values are also in reasonable agreement with published results from the same method [17, 38], there is a large variability among the resulting *K_A_*’s. For example, for PSM simulated at 48-55°C, Lee et al. [38] calculated a value of 456 ± 65 mN/m, which is similar to the one we obtained from the bilayer area fluctuation analysis (499 ± 42 mN/m), but Venable et al. [17] reported 310-350 mN/m, which is very similar to the moduli we obtained from NPγT simulations and with the LTF method. This variability seems more likely for high-melting lipids (PSM, DMPC, DPPC) whose dynamics 5-10°C above their melting temperatures are generally slower compared to low-melting temperature lipids, suggesting that the underlying reason for the divergent results is likely an insufficient sampling of the lateral area fluctuations. Indeed, in the analysis of box fluctuations each frame of the simulation trajectory represents a single data point, which makes proper sampling highly dependent on the length of the simulation and the size of the bilayer patch (the latter is closely related to the amplitude of the fluctuations [50]). To illustrate this point, we compare in Table 1 the *K_A_*’s for DOPC and DEPC (two fluid bilayers of the same size) calculated from either the full equilibrated trajectories of 517 and 680 ns respectively, or only from the last 100 ns of the simulation runs. The *K_A_* moduli obtained with bilayer area fluctuation analysis vary from 246 ± 20 mN/m to 313 ± 33 mN/m for DOPC and from 204 ± 18 mN/m to 321 ± 53 mN/m for DEPC, whereas those calculated with the LTF method do not show such variability. The reproducibility of the LTF moduli can also be seen from the analysis of replica simulations of DOPC, i.e. systems with identical size and lipid composition that were constructed and simulated independently from one another (Table 1).

For bilayers with up to 200 lipids, the results from our method are only weakly dependent on bilayer size: For POPC membranes with 128 and 200 lipids, we calculated *K_A_*’s of 245 ± 22 and 208 ± 20 mN/m respectively (Table 1). For a larger POPC bilayer with 416 lipids, the *K_A_* was 185 ± 18 mN/m, which may indicate a potential challenge for the analysis of larger systems, although the result is within the error of one of the smaller bilayers. Notably, since larger size of the simulated systems generally leads to larger undulations [50], the results might be affected by the assumption inherent in our method that the bilayer normal is the same throughout the bilayer surface (along the z axis of the simulation box). Because thicknesses are calculated by interpolation on the z positions of all atoms in the leaflet (see Methods), using the LTF method with the single bilayer normal assumption would tend to underestimate the apparent *K_A_* when large-scale undulations are present in the system. It is feasible to introduce local normals in the LTF formulation, but this would result in more complex numerical calculations. Instead, we sought to circumvent the problem by constraining the radius of interpolation, *R_int_* We reasoned that because large-scale bilayer undulations are known to appear on length scales larger than the bilayer thickness (*t_pp_*), setting *R_int_* to a length slightly larger than *t_pp_* could help alleviate the size dependence of *K_A_*. Indeed, using _(pè_ that is 10% larger than *t_pp_* produced a bilayer *K_A_* of 241 ± 23 mN/m for the larger POPC bilayer with 416 lipids, the same as calculated for the smaller systems.

Table 1 shows that the bilayer *K_A_* varied between 230 and 260 mN/m for most single-component bilayers we studied. This nearly constant *K_A_* across the bilayers is consistent with the idea that the bilayer compressibility is mostly influenced by the interfacial energy density, which arises from the common-to-all-bilayers hydrocarbon-water interactions at the leaflet-solvent interface [20]. In that respect, the somewhat higher compressibility of PSM (312 ± 29 mN/m) could be explained with the formation of intra- and intermolecular hydrogen bonds, characteristic for sphingomyelin molecules [51–53]. Interestingly, we found tetra-oleloyl cardiolipin (TOCL) to have a compressibility modulus of 231 ± 11 mN/m, in contrast to a previously reported modulus of 342 mN/m obtained from a rather poor linear fit of tension versus area expansion in a set of constrained tension simulations (Fig. 4 in Ref. [54]). The similarity of TOCL’s compressibility to that of a DOPG membrane (233 ± 19 mN/m) is particularly interesting because TOCL’s bending rigidity modulus was found to be twice that of DOPG [18]. (Note that chemically, TOCL resembles 2 DOPG lipids with linked headgroups).

**Figure 4.**
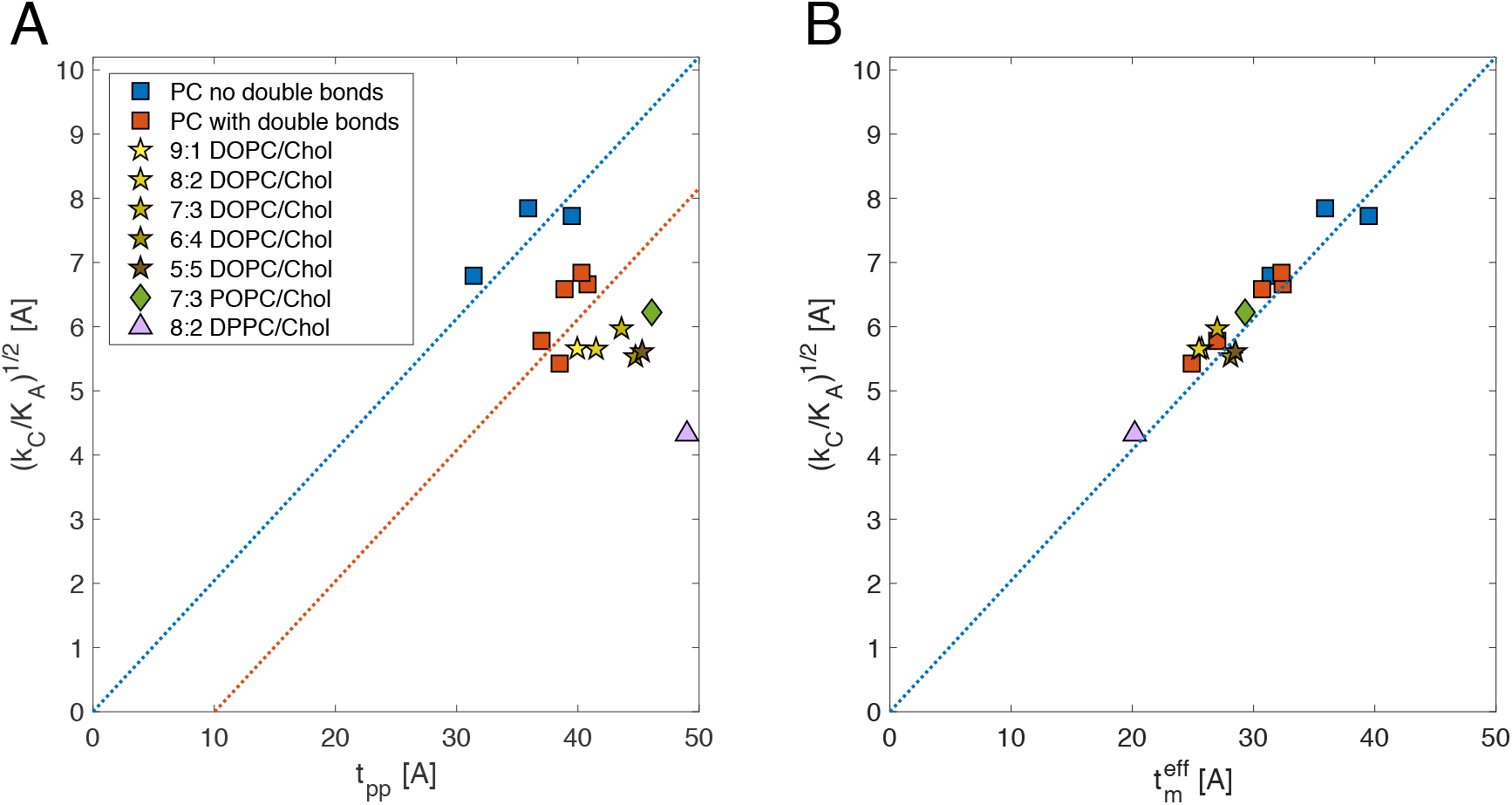
Application of the polymer brush model (PBM) to simulated bilayers suggests a new definition of bilayer mechanical thickness. A, The ratio of the bilayer bending rigidity (calculated from real-space analysis of splay fluctuations [18]) and area compressibility (calculated with the LTF method) from Eq. 17 is shown as a function of phosphate-to-phosphate distance, *t_pp_* for all single-component fully saturated (blue) and unsaturated (red) PC bilayers (Table 1) and binary mixtures of PC and Chol (yellow, Table 2). All dotted lines have been drawn with PBM’s slope of 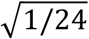, and different *x*-intercepts. A non-zero *x*-intercept indicates a deviation of the bilayer mechanical thickness, *t_m_*, from *t_pp_* as explained in the text. B, The ratio of the mechanical constants of the same bilayers in (A) are plotted against the *effective* mechanical thickness of the bilayers 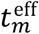, which is assumed to equal to: 1) *t_pp_* for all fully saturated PCs, 2) the difference between *t_pp_* and the length of the region around the double bonds for single-component unsaturated PCs and 9:1 DOPC/Chol, 3) the difference between *t_pp_* and the length of cholesterol’s ring body for all remaining binary mixtures of DOPC and Chol, and POPC and Chol, and 4) the difference between *t_pp_* and the full length of cholesterol for DPPC/Chol (Table S3). See text for more details.

### Application of the LTF method to mixed lipid systems

Interestingly, we found that binary mixtures of 30 mol% POPS with 70 mol% POPC or POPE at 20 and 25°C (respectively) have higher *K_A_*’s compared to most single-component bilayers. This is consistent with the combination of their large phosphate-to-phosphate thicknesses (40.1 and 42.8 Å) and small areas per lipid (60.9 and 55 Å^2^). Similarly, when increasing amounts of DMPC are added to POPC at 25°C, the bilayer *K_A_* increases gradually: from 210 ± 22 for 10 mol% DMPC to 262 ± 24 for 75 mol% DMPC, which is accompanied by a systematic decrease in the average bilayer thickness (from 38.6 to 36.6 Å) and area per lipid (from 63.8 to 61.4 Å2). Indeed, as shown in Fig. S4, we found a strong correlation (0.965) between the bilayer *K_A_* and the ratio of bilayer thickness to area per lipid for all fluid multi-component bilayers we studied, with the exception of the 8:2 DPPC/Chol membrane which is under gel-like conditions (see Table 2). Chol has a large effect on membrane compressibility (see Refs. in Table 2) and this is captured successfully by the LTF analysis as shown by the good agreement with experimental data (see Table 2 and Fig. 3). Indeed, the addition of Chol to DOPC at low concentrations (10 mol%) has a negligible effect on *K_A_*, but from approximately 20 mol% and higher, the *K_A_* value starts to increase gradually (see Fig. 3 and the discussion on the effect of Chol on bilayer mechanical thickness below). In agreement with results from experiments, we also find that Chol has a larger effect on the compressibility of POPC compared to DOPC: at 30 mol% Chol, POPC/Chol has a *K_A_* of 757 ± 87 mN/m while DOPC/Chol has a *K_A_* of 465 ± 57 mN/m. Interestingly, the large *K_A_* of the highly ordered 8:2 DPPC/Chol bilayer at 25°C (3126 ± 281 mN/m) is similar to the one reported for a 1:1 SM/Chol bilayer at 15°C (3327 ± 276 mN/m) [21].

### Ideal conditions for calculating area compressibility moduli with the LTF method

The accurate calculation of the area compressibility moduli relies on the equilibration of the bilayer area fluctuations. Thus, convergence of the bilayer lateral area must be ensured. One of the advantages of the LTF method is that, due to its local nature, it does not require long simulation times for analysis and also is not sensitive to system size. We recommend performing the analysis on at least 100-150 ns of equilibrated trajectory (or 5,000-10,000 frames), depending on bilayer size and thermodynamic state (e.g. smaller bilayers with a total of less than 150 lipids may need longer simulation times to obtain better statistics). If the chosen sampling is insufficient, this will result in larger errors on the *K_A_* that could be reduced by extending the simulation time.

Notably, in its current formulation, the LTF method assumes relatively small fluctuations in the membrane shape so that the local thicknesses can be obtained simply from the z-coordinates of the relevant groups of atoms (see section II.1). This assumption may not hold if the bilayer is large and develops strong undulations. In that case it may be necessary to specify a radius of interpolation for calculating local heights. Since the typical wavelength of such undulations appear on length scales larger than the bilayer thickness, this correction may need to be applied only on simulated bilayers whose thickness is much smaller than half the length of the simulation box (when utilizing periodic boundary conditions).

### Compression-bending relationship and the role of chain unsaturation

Another aspect of the validation of our method relates to the reproducibility of known relationships among the mechanical properties of the membrane. One basic principle of the mechanical properties of elastic sheets is that the compression (*K_A_*) and bending (*κ_C_*) moduli are related to one another through the sheet’s thickness *t_el_* up to some constant *C*, i.e., 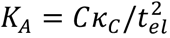. Since the elastic moduli of a lipid bilayer are calculated by assuming that the bilayer behaves as an elastic sheet, we tested whether the same compression-bending relationship holds for our systems. Using experimentally measured *K_A_*, *κ_C_* and phosphate-to-phosphate thickness of a number of phosphatidylcholine bilayers, this relation was initially demonstrated using *in vitro* data by treating each leaflet as a collection of freely jointed polymer chains [20]. The proposed simple model is referred to as the *polymer brush model* (PBM), which derives a proportionality constant *C* of 24 and thus gives:

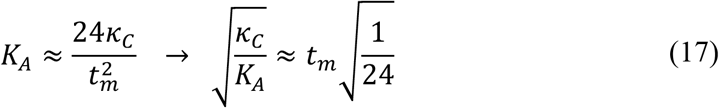

where *t_m_* is the *mechanical* thickness of the bilayer (i.e. the “deformable” thickness corresponding to *t_el_* in the above elastic sheet analogy). Since bilayer thickness is often measured as the average distance between the phosphate atoms of the two leaflets (*t_pp_*), *t_m_* can be expressed as *t_pp_* – *t_inc_* where *t_pp_* is some incompressible part within the length of the membrane. Notably, if *t_pp_* is the true mechanical thickness of the bilayer, then *t_inc_* = 0 and in a plot of *t_pp_* vs 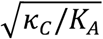, according to Eq. 17 above, all data points would lie on a line with slope 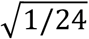 and *x*-intercept 0. However, *t_inc_* was found to be non-zero [20] and equal to 10 Å, implying that *t_m_* was 10 Å shorter than *t_pp_*. This difference was proposed to reflect the fact that the deformable thickness of a bilayer is limited to its hydrocarbon thickness, and since 5 Å represents the approximate vertical separation between the phosphorus atoms from the hydrocarbon acyl chains on either side of the bilayer, *t_m_* is 10 Å shorter than *t_pp_*, explaining the non-zero *t_inc_*. The only bilayers described as deviating from this behavior were polyunsaturated membranes [20], which appeared to have shorter mechanical thicknesses than expected.

We were able to test the relation between compression and bending and the applicability of PBM to the membrane systems we studied by taking advantage of the ability to calculate all three bilayer properties: bending rigidity [18], area compressibility and phosphate-to-phosphate thickness (Table S3) from our trajectories. Fig. 4A shows the results for all single-component PC bilayers together with the data for the binary mixtures of PC and Chol. Notably, the relationship between bending, compression and thickness could be explained with the PBM model (each dotted line has a slope of 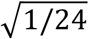 for all single-component bilayers. However, based on the results in Fig. 4A the membranes can be roughly divided into 2 categories with different mechanical thicknesses (*x*-intercepts): PC lipids with fully saturated acyl chains for which *t_m_* ≈ *t_pp_* (DLPC, DMPC, DPPC shown in blue) and lipids with 1 or more double bonds for which *t_m_* ≈ *t_pp_* − 10Å (POPC, SOPC, DEPC, DOPC, DLiPC shown in red in Fig. 4). We find that the data in Fig. 4A cannot be explained by the rationale for *t_inc_* given in Ref. [20], because the distance between the phosphates and the hydrocarbon chains on either side of the bilayer, proposed to be the origin of *t_inc_*, is independent of the saturation of the lipid chains (e.g. the average distance between the phosphates and the first carbons on the acyl chains is 4.7 Å for DPPC, 5.0 Å for POPC, 4.6 Å for DOPC and 4.6 Å for DLiPC). Instead, our result suggests an alternative model in which the double bonds are the ones responsible for reducing the deformable thickness.

To evaluate this alternative model, we tested if double bonds lead to relatively incompressible regions in the bilayer and decrease *t_m_* by increasing *t_pp_*. We first examined how pressure was distributed in tension-free bilayers with varying degrees of saturation (Fig. S5). An increase in the number of double bonds in the lipid chains lead to an increase in the pressure in the region around the double bonds. This observation prompted us to further investigate the relationship between pressure and compressibility in the context of lipid chain unsaturation. To do so, we calculated the *compressibility factor*, which is a correction factor in statistical mechanics that relates pressure, volume and number of particles (see below) and quantifies the deviation of the behavior (in particular, the compressibility) of a real gas from that of an ideal gas (or some other model matter with a defined equation of state) [55]. While the membrane is not an ideal gas (or a gas at all), the pressure at any given depth within the bilayer can be calculated in a slab with a defined volume that contains a certain number of atoms (see Section III in Methods), and hence the local state variables are well defined. Thus, we hypothesized that the double bonds may have an influence on compressibility that would be observable through this approximate model.

Following this model, we analyzed the pressure profile as a function of the average number of atoms in equal volume segments along the membrane normal. If the segments of the membrane each behaved like non-interacting ideal gasses, the pressure *P* can be expressed as a linear function of the number of atoms *n*, where the slope is proportional to the usual state variables:

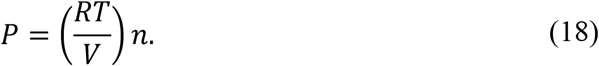

However, when pressure is plotted as a function of *n* for the saturated lipid bilayer DPPC, we find that *P* is inversely proportional to *n* (Fig. S6, row 1). As the relationship should not be negative, it is necessary to utilize an expanded state equation. The simplest expanded state equation is the van der Waals equation of state, which is often used to study fluids:

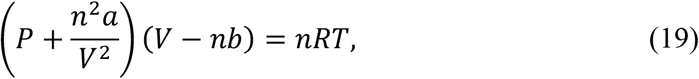

where *a* describes attractive interactions between atoms in the fluid and *b* describes the *excluded volume* (i.e. the volume occupied by the atoms that is excluded from the total volume) [55]. Eq. 19 can thus be rewritten as:

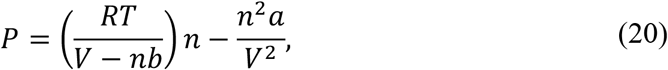

which reveals a strictly negative component that is a function of *n* and of the strength of the attractive interactions *a*. Thus, the negative relationship between *P* and *n* that is seen for DPPC can be explained by the inter- and intra-molecular bonds and interactions present in the bilayer. However, when analyzing membranes with varying degrees of unsaturation, we find that upon adding double bonds, a regime of positive slope occurs (Fig. S6, rows 2-4). This observation can be explained by the dominance of the first term in Eq. 20 due to the increase in excluded volume caused by the presence of double bonds, which limit the space that can be sampled by the lipid chains. Indeed, a comparison between the van der Waal’s constants for ethylene (C_2_H_4_ *b* = 0.05821 L/mol), ethane (C_2_H_6_ *b* = 0.06499 L/mol) and hydrogen (H_2_ *b* = 0.02651 L/mol) [55, 56] clearly shows that a double bond increases the *b* term (i.e. 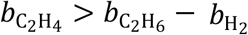). Further comparison of the corresponding *a* terms (C_2_H_4_ *a* = 4.612 bar L^2^/mol^2^, C_2_H_6_ *a* = 5.570 bar L^2^/mol^2^, H_2_ *a* = 0.2453 bar L^2^/mol^2^) suggests that a double bond also decreases the attractive interactions between atoms (i.e. 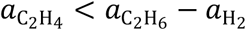), which would further contribute to the dominance of the first term in Eq. 20.

To investigate the expected change in compressibility due to the additional double bonds we interpreted the data in Fig. S5 in the context of the compressibility factor. When quantifying the deviation from an ideal gas, the compressibility factor is defined as:

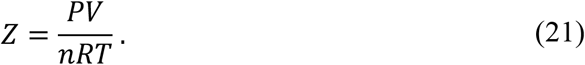

When *Z* > 1, the volume is greater than expected for a given pressure due to its incompressibility. When combined with Eq. 20, the compressibility factor for a van der Waals fluid becomes a function of the excluded volume and the interaction between atoms,

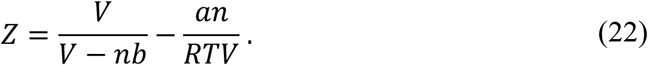

In the regime in which the excluded volume term dominates (i.e. when there are double bonds present), Z becomes large and positive, indicating the membrane is less compressible. This analysis thus confirms that the addition of double bonds to lipid chains leads to relatively incompressible regions that decrease *t_m_* by increasing *t_inc_*.

Motivated by the effect of the double bonds on the chain order parameter profile of the lipids, we used a simple heuristic approach to approximate the length of the perturbed region around the double bonds, *t_DB_*, for the unsaturated PC bilayers from Fig. 4A: namely, the perturbed region extends 4 carbons above and below the midpoint of all double bonds for DOPC and DLiPC, and 2 carbons above and below the midpoint of the double bonds for POPC, SOPC and DEPC (see Fig. S7 and Table S3). Remarkably, after subtracting *t_DB_* from *t_pp_* and replotting the data in Fig. 4B, all data points moved to the line with *x*-intercept at 0, confirming our hypothesis regarding the nature and source of the incompressibility, and suggesting that the bilayer mechanical thickness for unsaturated lipids can be defined as the phosphate-to-phosphate thickness *excluding* the regions around the double bonds. We note that this result explains as well the observation in Ref. [20] that polyunsaturated bilayers have shorter mechanical thicknesses, since the perturbed region around their double bonds would be larger and consequently, *t_m_* would be smaller.

Although DLiPC (di18:2 PC) has two more double bonds per molecule than DOPC (di18:1 PC), we find that both DLiPC and DOPC bilayers have similar mechanical thicknesses. This result is consistent with the nearly identical chain stress distributions in monolayers of these two lipids, predicted by Cantor [57]. As illustrated by the analysis in Ref. [57], the effect of the double bonds on monolayer properties (and very likely, bilayer properties) depends on both the number and the location of the double bonds.

Note that in the validation of the PBM described in Ref. [20], the mechanical thickness of DMPC was found to be 10 Å shorter than *t_pp_*, like the rest of the examined bilayers. However, the reported DMPC’s bending rigidity modulus measured by micropipette aspiration (13.2 kT) was much lower than the one obtained with flicker spectroscopy (31.1 kT, [58]) and also with our computational method (34.7 kT, [18]). The low value of *κ_C_* reported in Ref. [20] is most likely due to the difficulties of applying the micropipette aspiration technique to higher melting temperature lipids as discussed in Ref. [18]. If a higher *κ_C_* had been used instead in the PBM analysis, DMPC’s mechanical thickness would likely have been predicted to be much closer to *t_pp_*, consistent with the data in Fig. 4.

### The effect of cholesterol on bilayer mechanical thickness

As illustrated by the analysis above, the effective mechanical thickness of a bilayer, 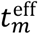, can be expressed as the phosphate-to-phosphate distance excluding the length of any region that resists compression. In this respect, it is interesting to investigate the behavior of lipid bilayers containing cholesterol, as the sterol contains a rigid and relatively incompressible set of rings [47]. Fig. 4A shows the mechanical properties of various lipid mixtures with cholesterol, including a set of DOPC/Chol bilayers with varying amounts of Chol tested against PBM. Most of these bilayers exhibit a larger deviation from either of the dotted lines, indicating a larger *t_inc_* than the non-Chol membranes. To determine *t_inc_* for the DOPC/Chol and POPC/Chol membranes, we first used the same heuristic approach as for the unsaturated lipids described above and calculated 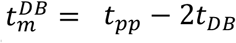. With this definition, all DOPC/Chol data points from Fig. 4A shifted to the line with an x-intercept at 0, but for the POPC/Chol bilayer 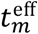 was shorter than 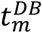 (Fig. S8A). Given the larger effect of Chol on POPC compared to DOPC and the relatively incompressible and rigid nature of the sterol ring structure as discussed above, we reasoned that in the POPC/Chol membrane *t_inc_* would be better approximated by the length of cholesterol’s ring region, *t_Chol_*. To test this hypothesis, we calculated *t_Chol_* as the average distance between the C3 and C17 atoms of Chol (using CHARMM36 notation) projected onto the *z*-axis, and found it to be 8.4 Å (Table S3). Remarkably, plotted as a function of 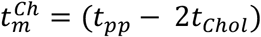, the PBM data-point for POPC/Chol fell on the line with x-intercept at 0 (Fig. S8B), indicating that 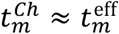.

We then tested how well 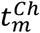 approximates 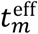 in the DOPC/Chol mixtures. In light of the results in Fig. 3 showing that Chol has an effect on the bilayer *K_A_* only at 20 mol% or higher, we expected that *t_inc_* ≈ 2*t_Chol_* for these Chol bilayers but not for the 9:1 DOPC/Chol membrane. Indeed, as shown in Fig. S8B, 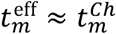 for the 20-50% DOPC/Chol mixtures, and 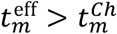) for the 10% DOPC/Chol bilayer.

Three different regimes are required to describe the effect of Chol on the structural properties of bilayers, corresponding to low, intermediate, and high cholesterol mole fractions [47, 59]. The results presented above and summarized in Figs. 3 and 4 suggest that this is also the case for the Chol effects on bilayer mechanical thickness and compressibility. Thus, our analysis shows that at 10 mol%, Chol does not have an effect on either 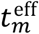 or *K_A_*; at 20 and 30 mol% Chol affects 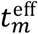 and increases *K_A_ indirectly* through its condensing effect on DOPC; and at 40 and 50 mol% Chol affects 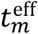 and increases *K_A_* directly, i.e. its contribution to *K_A_* must be considered explicitly (Fig. 3). These regimes are consistent with the reduction in the translational and rotational entropy for Chol with increasing concentration: at low mole fractions Chol adopts relatively large tilt angles with respect to the bilayer normal, and thus relatively random orientations (large tilt angles result in larger degeneracy of rotational states [18, 60]). This increased orientational freedom can alleviate any potential stress from compressing the bilayer’s thickness. At higher concentrations, Chol’s conformational freedom gradually decreases, as Chol molecules tend to align parallel to the bilayer normal [60]. Bilayer compression under such conditions likely involves compression of the Chol molecules.

We also investigated the applicability of the PBM to a fully saturated DPPC/Chol bilayer with 20 mol% Chol. The bilayer was simulated at 25°C and is in a very ordered gel-like state as indicated by its small area per lipid and large elastic moduli (Tables 2, S1, S3). The PBM comparison for this system, shown in Fig. 4A, indicates that the mechanical thickness of the membrane is significantly smaller than *t_pp_*. Interestingly, we found that for this bilayer the Chol tail was more rigid than in the other membranes (Fig. S9A), and comparable in its tilt distribution to the ring body of Chol in the 7:3 DOPC/Chol bilayer (Fig. S9B), suggesting that it was harder to compress, and that the mechanical thickness of the membrane could exclude the sterol tail as well due to the high order in the system. Indeed, considering in the calculations the full length of Chol, including its ring and tail regions, we were able to successfully recover 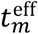 of the DPPC/Chol bilayer (Fig. 4B). Both Chol concentration and the temperature determine the thermodynamic phase behavior of the bilayer and thereby the degree of Chol’s conformational freedom. Therefore, it remains to be investigated whether Chol affects the mechanical properties of other fully saturated lipids in a similar way, and how the observed relationships vary with temperature.

### Revised definition of bilayer mechanical thickness clarifies conflicting reports on PBM’s applicability

As illustrated by our analysis, the effect of double bonds and cholesterol on bilayer’s mechanical thickness has not been systematically characterized before. Therefore, we sought to examine the relation of our findings to published observations from both *in vitro* and *in silico* work. We found that our results resolve some contradictory reports in the literature regarding the validity of PBM for different bilayers. Indeed, since PBM was first introduced, the model has been experimentally tested in a number of studies by assuming that *t_m_* = *t_pp_* − 10 Å, yielding conflicting results [17, 27, 28]. In 2008 Pan et al. found a good agreement between their results and PBM’s predictions for DOPC [27] but in 2009 they reported that the theory was incapable of describing the relationship between the mechanical constants in cholesterol-containing bilayers [28]. In a comprehensive review of bilayer mechanical properties from MD simulations published more recently, Venable et al. found a relatively good agreement between the simulation results and PBM for POPC and DOPC but there was a bigger deviation for DMPC and DPPC (see Fig. 11 in Ref. [17]). These conflicting observations can be consolidated in light of our finding that the presence of cholesterol and/or acyl chain unsaturation can affect the mechanical thickness of the bilayers. Not surprisingly, PBM was successfully applied to lipids with 1 or 2 double bonded tails such as DOPC and POPC both in the experimental and computational studies, since for those lipids *t_m_* can indeed be approximated by *t_pp_* − 10 Å. However, according to our analysis, the incompressible body of cholesterol has length of about 8.3 Å, effectively decreasing *t_m_* by an additional 6 Å (3 Å from each leaflet), i.e. *t_m_* ≈ *t_pp_* − 16 Å. Thus, if the hydrocarbon thickness 2*D_C_* (used as a proxy to *t_pp_* − 10 Å) in Eq. 6 in Ref. [28] is substituted with (2*D_C_* − 6 Å), the ratio between predicted and actual *κ_C_* for SOPC and DOPC in the presence of 30 mol% cholesterol becomes 1.0 and 1.36, respectively, indicating a good agreement with the theory contrary to the stated conclusion in Ref. [50]. Similarly, since DMPC and DPPC are fully saturated and hence *t_m_* ≈ *t_pp_*, the use of *t_pp_* − 10 Å for their mechanical thickness in Ref. [17] explains the larger deviation of their calculated bending moduli from PBM’s predictions.

Together, all these results are consistent with the notion that bilayer mechanical thickness depends on lipid composition and cannot be simply taken as the bilayer hydrocarbon thickness. Our analysis illustrates a simple but general principle of bilayer mechanics whereas the assumption for elastic material behavior holds only for the regions within the membrane that are equally compressible. In particular, the presence of both acyl chain unsaturation and cholesterol produce non-uniform compressibility in the membrane hydrocarbon core that needs to be taken into account when quantifying the deformable membrane thickness.

## CONCLUSION AND SUMMARY

We have presented a new computational framework for calculating area compressibility moduli of lipid bilayers and their individual leaflets from all-atom MD simulations. The approach is based on sampling local thickness fluctuations from an MD simulation trajectory in which these properties are sufficiently converged. We show that the method overcomes a number of limitations of existing computational approaches, and yields elastic moduli values that are in agreement with available experimental data for both single and multi-component bilayers composed of saturated and unsaturated lipids, and cholesterol, simulated at several temperatures. Importantly, because it is free from the need to sample global lateral bilayer fluctuations, our method is uniquely capable of analyzing the area compressibility of bilayers under tension (i.e. simulated in an NPAT ensemble). We note that it should also allow, in principle, future applications for the calculation of the spatial variability in leaflet compressibility moduli in the presence of transmembrane inclusions.

The data presented show that the mechanical properties of the simulated bilayers, and the relation between specific parameters representing their properties, are consistent with an elastic sheet model and consonant with a polymer brush model (PBM). However, the application of the PBM is shown to require a significant modification of the canonical definition of the membrane mechanical thickness considered as simply the hydrocarbon region of the bilayer. Indeed, we demonstrated the specific considerations that are necessary to determine the appropriate mechanical thickness required to calculate unknown elastic moduli. These include the quantitative accounting for acyl chain unsaturation and cholesterol concentration, both of which introduce relatively incompressible regions within the bilayer that decrease the effective mechanical thickness.

While all of the bilayers we have examined have the same lipid composition and the same number of lipids in their two leaflets, the physical model underlying Eq. 10 is general enough to allow for analysis of the great variety of compositionally asymmetric bilayers that are physiologically relevant, as their local leaflet *K_A_*-s could be different. This aspect of the method continues to be the subject of our ongoing computational studies. Systematic experimental measurements of the compressibility moduli of asymmetric bilayers, especially ones whose leaflets are expected to have significantly different mechanical properties, would greatly benefit the validation and/or refinement of the theoretical predictions from the harmonic mean relationship between the bilayer *K_A_* and the local leaflet moduli (Eq. 10).

## Author Contributions

MD, GK, ML and HW designed the research and wrote the manuscript. MD performed all simulations and computational analysis.

## ACKNOWLEDGMENT

We thank Daniel Harries and Alex Sodt for insightful discussions, and Fred Heberle for valuable help with the analysis related to testing the volume incompressibility assumption. The work was supported by NIH Grant P01 DA012408 and NSF (Award NSF 1740990). The extensive computational work used the following resources: the Extreme Science and Engineering Discovery Environment (XSEDE, accounts TG-MCB120008 and TG-MCB130010), which is supported by National Science Foundation grant number ACI-1053575; an allocation at the National Energy Research Scientific Computing Center (NERSC, repository m1710) supported by the Office of Science of the U.S. Department of Energy under Contract No. DE-AC02-05CH11231; and the computational resources of the David A. Cofrin Center for Biomedical Information in the HRH Prince Alwaleed Bin Talal Bin Abdulaziz Alsaud Institute for Computational Biomedicine at Weill Cornell Medical College.

## SUPPORTING MATERIAL

**Table S1.** Bilayer information for all simulated systems: number of lipids per leaflet (Lip/Leaf), temperature (Temp), salt concentration, number of water molecules per lipid (N H_2_O), trajectory length (both total and analyzed) and average area per lipid (APL). Also shown are the standard errors for the area per lipid, calculated from consecutive time blocks of length determined by the effective number of uncorrelated data points [1].

| Bilayer | Lip/Leaf | Temp<br>[°C] | Salt<br>[2] | N<br>H<sub>2</sub>O | Traj. Length<br>[ns] |  | APL [Å<sup>2</sup>] |
| --- | --- | --- | --- | --- | --- | --- | --- |
|  |  |  |  |  | analysis | total |  |
| DLPC* | 64 | 30 |  | 45 | 278 | 278 | 62.6 ± 0.1 |
| DMPC* | 64 | 30 | 140 | 50 | 200 | 200 | 60.4 ± 0.1 |
| DPPC* | 100 | 50 |  | 45 | 238 | 335 | 61.5 ± 0.1 |
| DLiPC* | 100 | 25 |  | 45 | 245 | 278 | 70.1 ± 0.1 |
| DOPC* | 100 | 25 |  | 45 | 517 | 523 | 68.2 ± 0.1 |
| DOPC replica 1 | 100 | 25 |  | 45 | 269 | 275 | 68.1 ± 0.1 |
| DOPC replica 2 | 100 | 25 |  | 45 | 267 | 274 | 68.1 ± 0.1 |
| SOPC* | 100 | 25 |  | 45 | 464 | 490 | 63.8 ± 0.1 |
| DEPC* | 100 | 25 |  | 45 | 680 | 680 | 64.4 ± 0.1 |
| POPC* 1 | 208 | 25 |  | 45 | 520 | 520 | 64.4 ± 0.1 |
| POPC* 2 | 100 | 25 | 140 | 70 | 183 | 226 | 64.3 ± 0.2 |
| POPC 3 | 64 | 25 |  | 45 | 281 | 281 | 64.3 ± 0.1 |
| POPC 4 | 64 | 30 |  | 45 | 293 | 293 | 64.7 ± 0.1 |
| POPE* | 100 | 55 |  | 45 | 184 | 190 | 60.8 ± 0.1 |
| DOPG* | 100 | 25 | 140 | 60 | 231 | 246 | 71.2 ± 0.1 |
| TOCL* | 50 | 30 | 140 | 60 | 310 | 310 | 130.5 ± 0.2 |
| PSM | 100 | 55 |  | 45 | 235 | 324 | 56.3 ± 0.1 |
| SAPE | 100 | 37 |  | 45 | 563 | 928 | 63.6 ± 0.1 |
| POPC/Chol 70/30 | 100 | 15 |  | 45 | 252 | 1157 | 45.1* ± 0.1 |
| DOPC/Chol 90/10 | 100 | 25 |  | 45 | 336 | 336 | 62.3* ± 0.1 |
| DOPC/Chol 80/20 | 100 | 25 |  | 45 | 350 | 350 | 56.9* ± 0.1 |
| DOPC/Chol 70/30 | 100 | 15 |  | 45 | 329 | 676 | 50.8* ± 0.1 |
| DOPC/Chol 60/40 | 100 | 25 |  | 45 | 179 | 465 | 47.1* ± 0.1 |
| DOPC/Chol 50/50 | 100 | 15 |  | 45 | 859 | 948 | 43.8* ± 0.0 |
| DPPC/Chol 80/20* | 100 | 25 |  | 45 | 124 | 683 | 40.8* ± 0.0 |
| POPE/POPG 70/30* | 100 | 37 |  | 60 | 274 | 280 | 61.2 ± 0.1 |
| POPC/POPS 70/30* | 70 | 20 | 50 | 45 | 175 | 191 | 60.9 ± 0.1 |
| POPE/POPS 70/30* | 210 | 25 |  | 82 | 250 | 690 | 55.3 ± 0.1 |
| DMPC/POPC 10/90 | 100 | 25 |  | 45 | 471 | 471 | 63.8 ± 0.1 |
| DMPC/POPC 43/57 | 100 | 25 |  | 45 | 556 | 696 | 62.6 ± 0.1 |
| DMPC/POPC 75/25 | 100 | 25 |  | 45 | 738 | 738 | 61.4 ± 0.1 |

\* Bilayers taken from Ref [3].
\* The given APLs include cholesterol.

**Table S2.** Information for all NPγT simulations. For each bilayer shown are the set of applied non-zero tensions (*γ*), the corresponding average areas per lipid (APL) and lengths of the last part of the trajectories used for analysis, as identified by the method from Ref. [1] (see Simulations analysis and method implementation section in the text).

| Bilayer | $\gamma$ [mN/m] | APL [$\text{\AA}^2$] | Lengths of analyzed trajectories [ns] |
| --- | --- | --- | --- |
| DMPC | -5, 3, 5, 12 | 58.7, 61.0, 61.6, 63.2 | 168, 314, 100, 302 |
| DPPC | -3, 5, 9 | 60.8, 62.8, 64.1 | 267, 239, 244 |
| DLiPC | -3, 4, 8, 12 | 69.5, 71.0, 72.4, 73.9 | 172, 173, 180, 183 |
| DOPC | -7, 7, 15 | 66.3, 70.2, 72.2 | 138, 144, 129 |
| POPC | -5, 4, 8 | 62.8, 65.1, 66.8 | 91, 191, 234 |
| POPE | -3, 3, 5, 9 | 60.1, 61.3, 61.9, 62.7 | 198, 196, 193, 168 |
| PSM | -3, 3, 10 | 55.6, 56.7, 57.9 | 342, 221, 128 |

**Table S3.** Bilayers used for comparison with PBM (Fig. 4). Shown are the average phosphate-to-phosphate distance (*t_pp_*), length of the double bond region approximated as described in the caption of Fig. S7 (*t_DB_*), length of cholesterol’s ring body 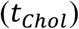, the difference between *t_pp_* and 2*t_DB_* 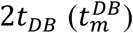 or *t_pp_* and 2*t_Chol_* 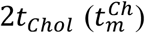, the effective mechanical thickness of the bilayers 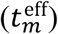, and bilayer bending rigidity (*κ*_C_) calculated from splay fluctuations [4–6]. Errors for all thicknesses shown in parenthesis are the standard deviations from consecutive time blocks of length determined by the effective number of samples [1]. Errors on *κ*_C_ were calculated as described in Ref. [6]. In the first column, DOPC/Chol, POPC/Chol and DPPC/Chol mixtures are denoted with ‘DO’, ‘PO’ and ‘DP’ respectively.

| Bilayer | $t_{pp}$ [$\text{\AA}$] | $t_{DB}$ [$\text{\AA}$] | $t_{chol}$ [$\text{\AA}$] | $t_m^{DB}$ [$\text{\AA}$] | $t_m^{Ch}$ [$\text{\AA}$] | $t_m^{\text{eff}}$ [$\text{\AA}$] | $\kappa_C$ [$k_B T$] |
| --- | --- | --- | --- | --- | --- | --- | --- |
| DLPC | 31.4 (0.4) |  |  |  |  | 31.4 (0.4) | 25.8 (0.6)* |
| DMPC | 35.9 (0.4) |  |  |  |  | 35.9 (0.4) | 34.7 (1.2)* |
| DPPC | 39.5 (0.4) |  |  |  |  | 39.5 (0.4) | 34.1 (1.1)* |
| DLiPC | 37.0 (0.3) | 5.0 (0.1) |  | 27.0 (0.4) |  | 27.0 (0.4) | 16.3 (0.3)* |
| DOPC | 38.5 (0.3) | 6.8 (0.1) |  | 24.9 (0.4) |  | 24.9 (0.4) | 18.3 (0.3)* |
| SOPC | 40.8 (0.4) | 4.2 (0.1) |  | 32.4 (0.4) |  | 32.4 (0.4) | 26.4 (0.7)* |
| DEPC | 40.3 (0.4) | 4.0 (0.1) |  | 32.3 (0.4) |  | 32.3 (0.4) | 24.2 (0.6)* |
| POPC | 38.9 (0.4) | 4.1 (0.1) |  | 30.7 (0.4) |  | 30.7 (0.4) | 24.3 (0.6)* |
| PO 7:3 | 46.1 (0.3) | 5.3 (0.1) | 8.4 (0.1) | 35.5 (0.4) | 29.3 (0.4) | 29.3 (0.4) | 73.7 (1.3) |
| DO 9:1 | 39.9 (0.4) | 7.1 (0.1) | 7.9 (0.2) | 25.7 (0.4) | 24.1 (0.6) | 25.7 (0.4) | 22.5 (0.4) |
| DO 8:2 | 41.5 (0.4) | 7.6 (0.1) | 8.0 (0.1) | 26.3 (0.4) | 25.5 (0.4) | 25.5 (0.4) | 30.5 (0.6) |
| DO 7:3 | 43.4 (0.4) | 8.2 (0.1) | 8.2 (0.1) | 27.0 (0.4) | 27.0 (0.4) | 27.0 (0.4) | 41.6 (0.7) |
| DO 6:4 | 44.7 (0.3) | 8.7 (0.1) | 8.3 (0.1) | 27.3 (0.4) | 28.1 (0.4) | 28.1 (0.4) | 51.0 (0.6) |
| DO 5:5 | 45.3 (0.3) | 9.0 (0.1) | 8.4 (0.0) | 27.3 (0.4) | 28.5 (0.3) | 28.5 (0.3) | 70.1 (1.1) |
| DP 8:2 | 49.0 (0.2) |  | 14.4 (0.1)* |  | 20.2 (0.4) | 20.2 (0.4) | 142.1 (3.1)* |

\* Taken from Ref [3].
\* $t_{chol}$ includes both the ring (8.4 $\text{\AA}$ ) and tail (6 $\text{\AA}$ ) regions of cholesterol.

**Figure S1.**
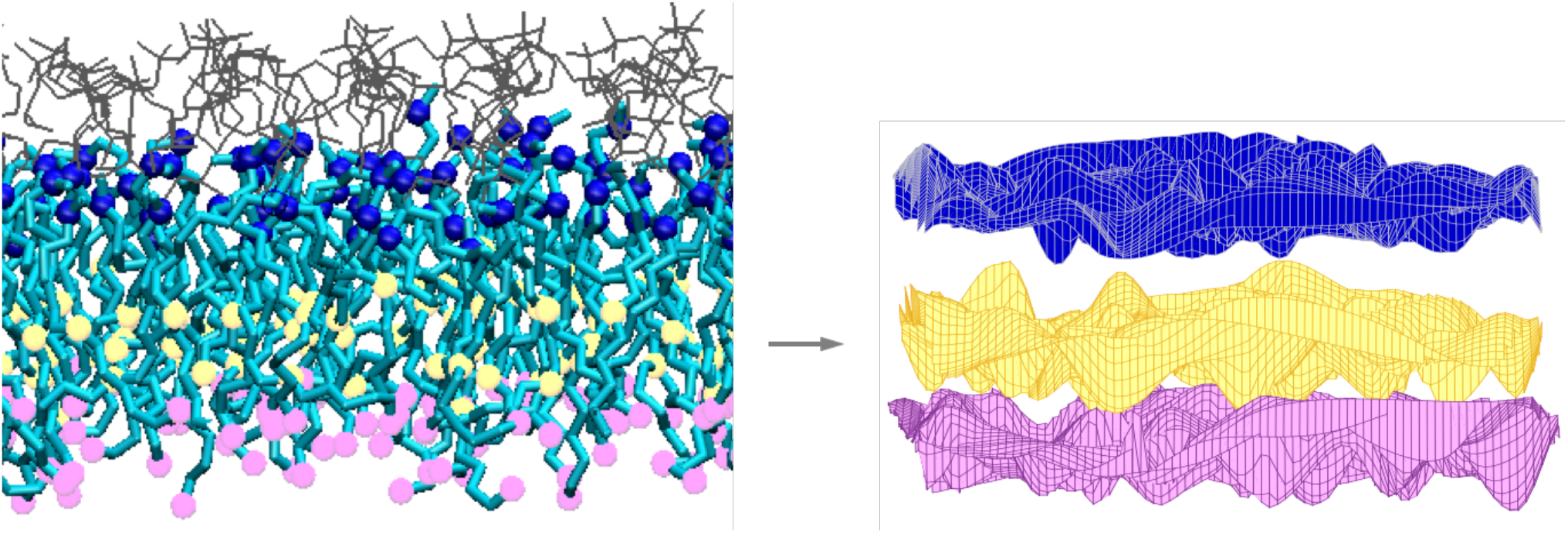
Transformation of a leaflet from an all-atom MD simulation trajectory frame into a corresponding set of surfaces. Left, a schematic representation of a DPPC leaflet. Lipid chains are shown in Licorice (cyan), the first carbons not attached to oxygen are shown as blue spheres, the 10th carbons (representing the surface for the relevant thickness fluctuations for this membrane, see text) are shown as yellow spheres and the terminal methyl carbons are shown as pink spheres. The glycerol backbone atoms, phosphates and lipid headgroups are shown as grey lines. Right, three interpolated surfaces corresponding to the blue, yellow and pink spheres on the left.

**Figure S2.**
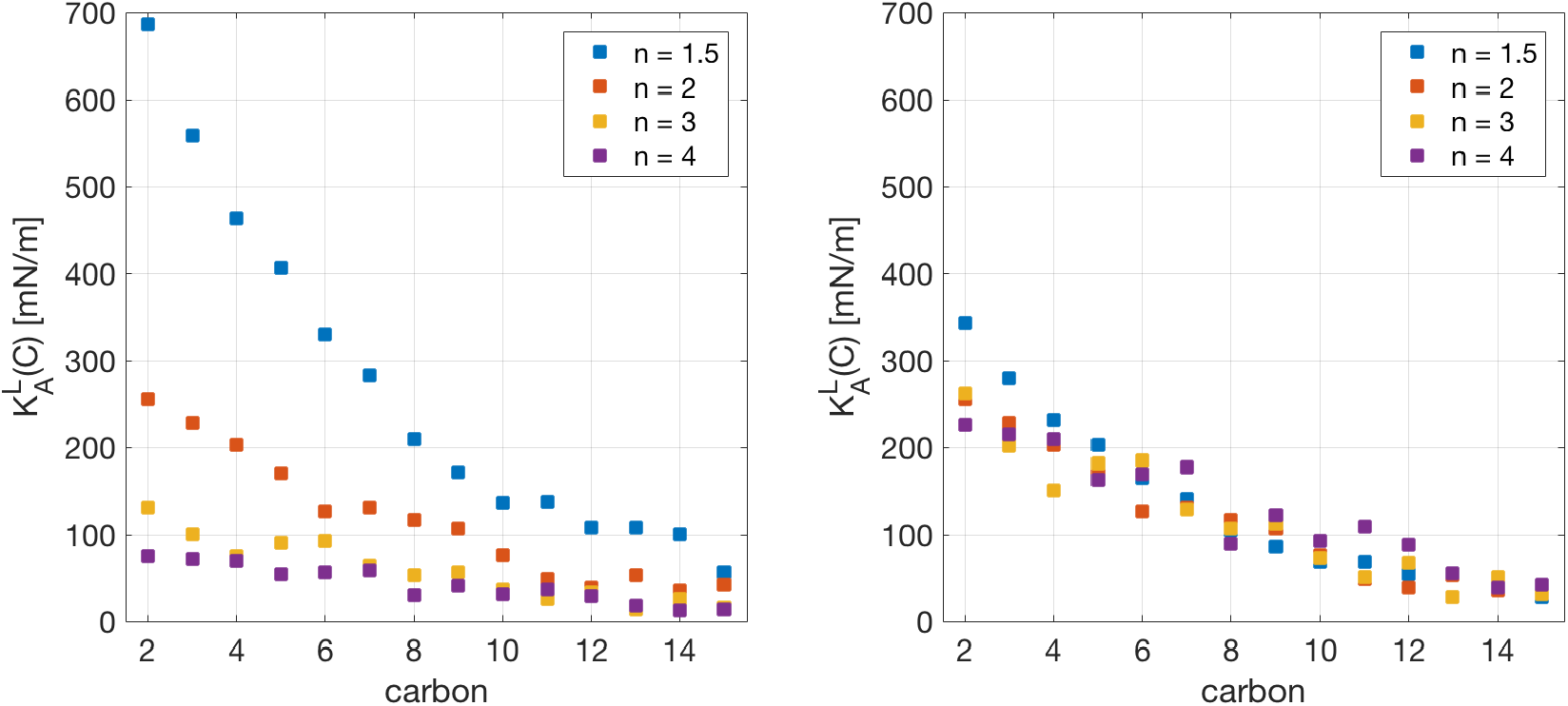
Effect of the interpolation order *n* on the calculated effective leaflet compressibilities (y-axis) at different carbon surfaces (x-axis) by setting *a*_3_ to either 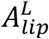 (left) or *a*_3,8_ (right). The data shown are for the top leaflet of a DOPC bilayer.

**Figure S3.**
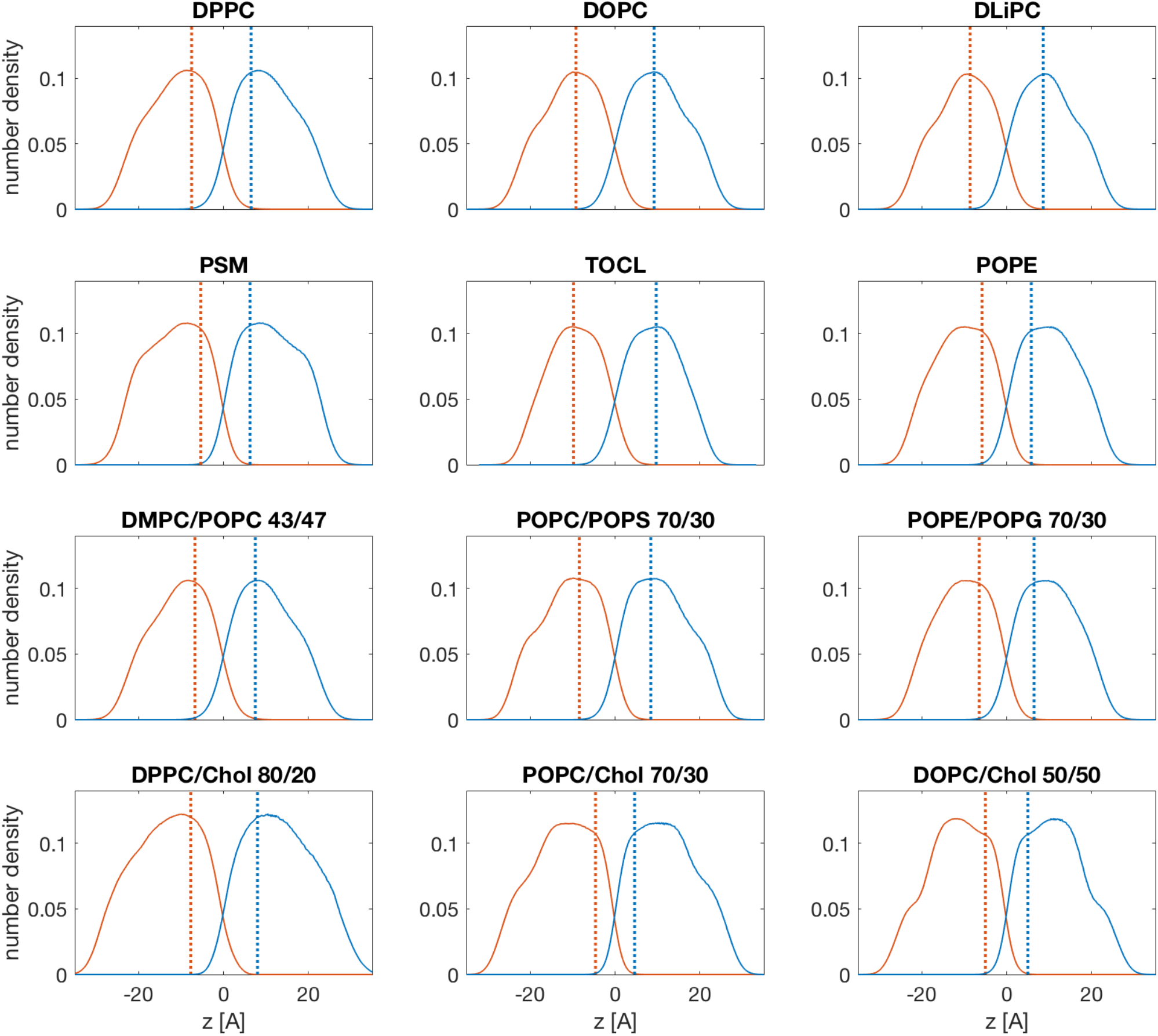
Identified leaflet thicknesses whose fluctuations are relevant for the calculation of the leaflet compressibility moduli for a set of representative bilayers from Tables 1–2. Shown are the number density profiles of the top (blue) and bottom (red) bilayer leaflets. The corresponding locations of the identified relevant leaflet thicknesses are shown as dotted lines.

**Figure S4.**
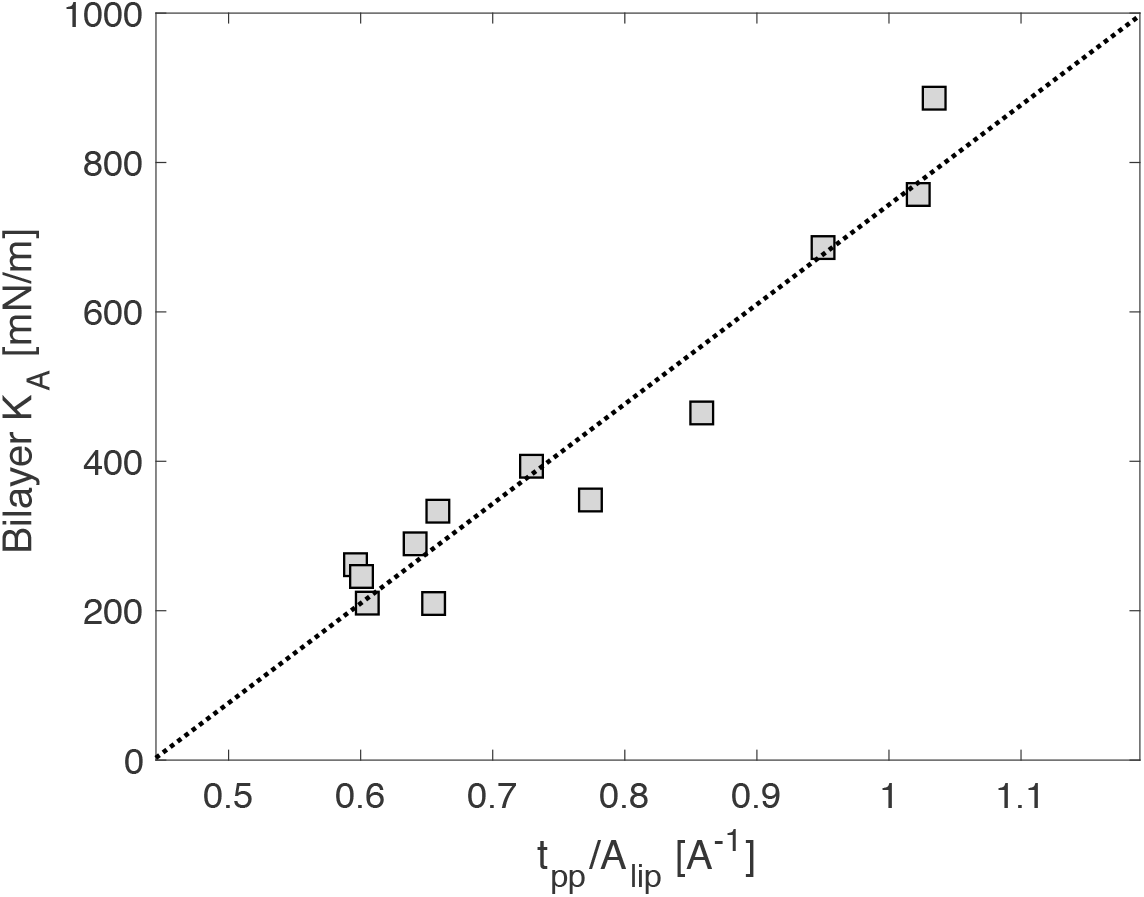
Bilayer area compressibility modulus *K_A_* as a function of the ratio of bilayer thickness (phosphate-to-phosphate distance, *t_pp_*) and average area per lipid (*A_lip_*) for all fluid multi-component bilayers from Table 2, except for DPPC/Chol. The correlation between *K_A_* and *t_pp_*/*A_lip_* is 0.965. The best linear fit to the data is shown as a dotted black line.

**Figure S5.**
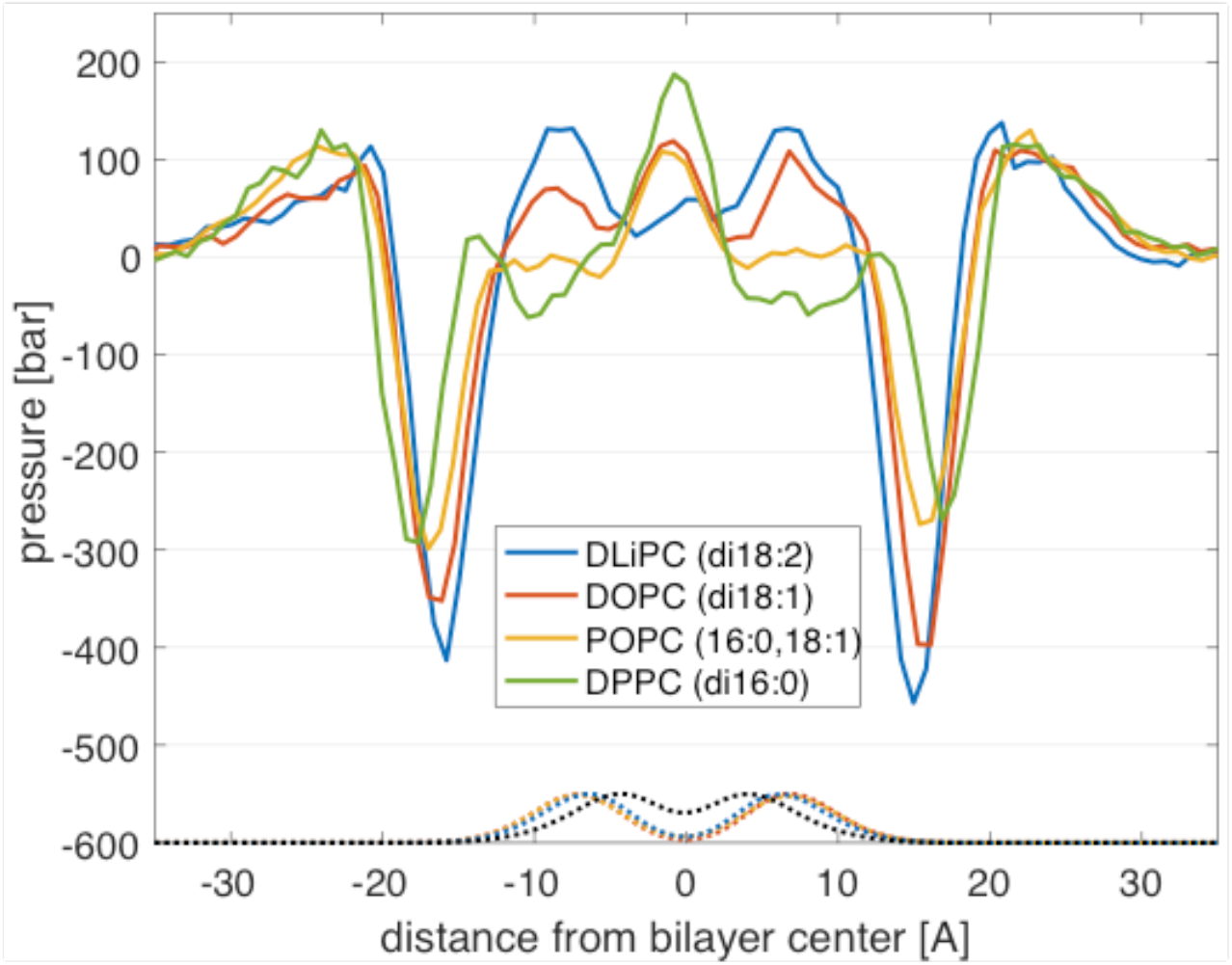
Lateral pressure profiles of bilayers with increasing level of unsaturation. All double bonds are between carbons 9-10, or 9-10 and 12-13 for DLiPC. An increase in pressure is observed with an increase in the number of double bonds (see text for more details). Also shown with correspondingly colored dotted lines are the rescaled and translated number density profiles of the double bonds at carbons 9-10 for all unsaturated bilayers. The number density of the second double bond at carbons 12-13 for DLiPC is shown in black. The pressure profiles have been smoothed for visualization purposes.

**Figure S6.**
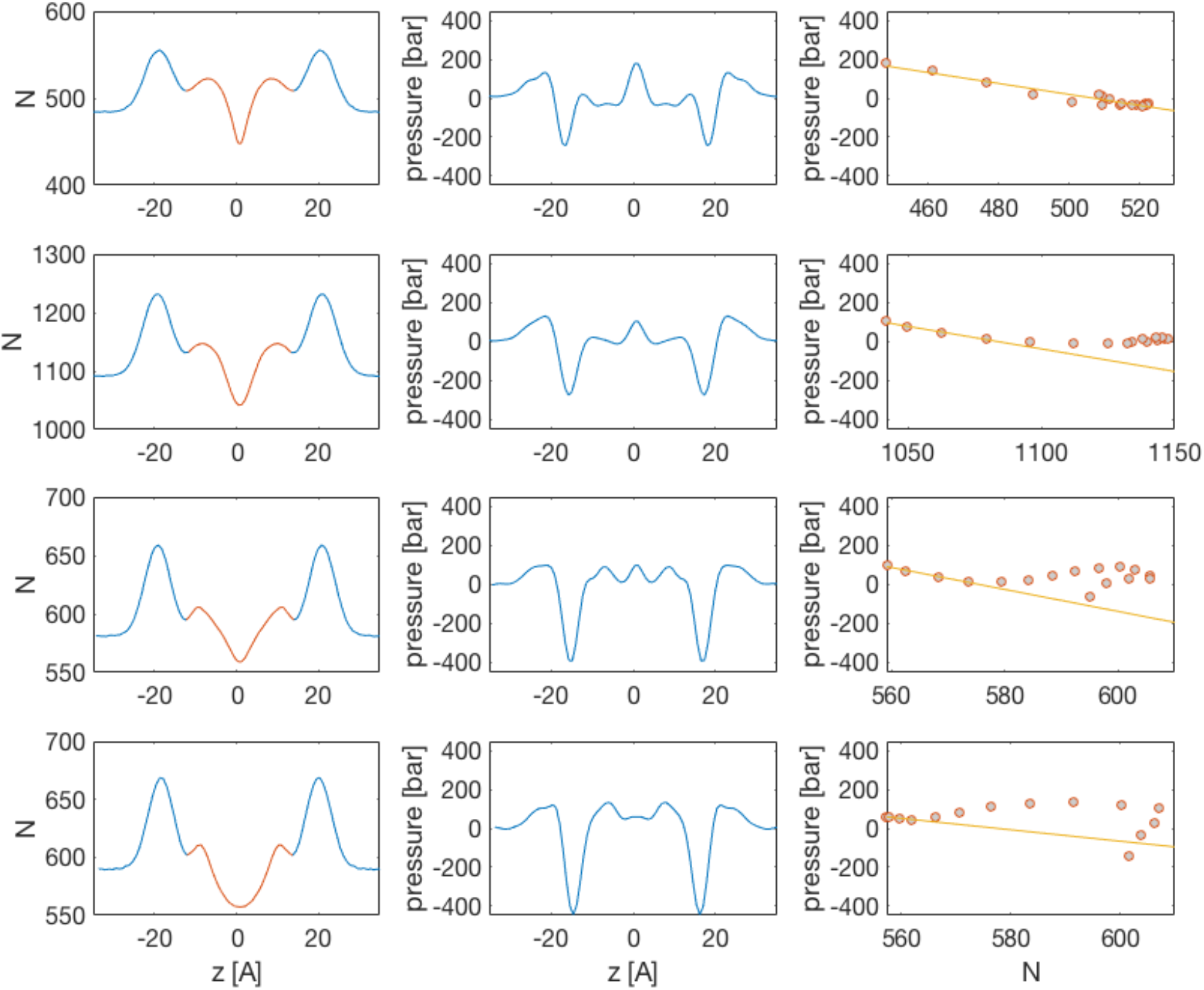
Total number of atoms N (**first column**) and lateral pressure p (**second column**) calculated from the same equal volume slabs along the bilayer normal for bilayer with increasing number of double bonds: DPPC (first row), POPC (second row), DOPC (third row) and DLiPC (fourth row). The **third column** shows the corresponding relationship between N and p in the central region (colored in red) in the N vs z plots from the first column. A best-fit line through the left-most points on the plots in the third column (corresponding to the slabs around the bilayer midplane at z=0) is shown in yellow to guide the eye.

**Figure S7.**
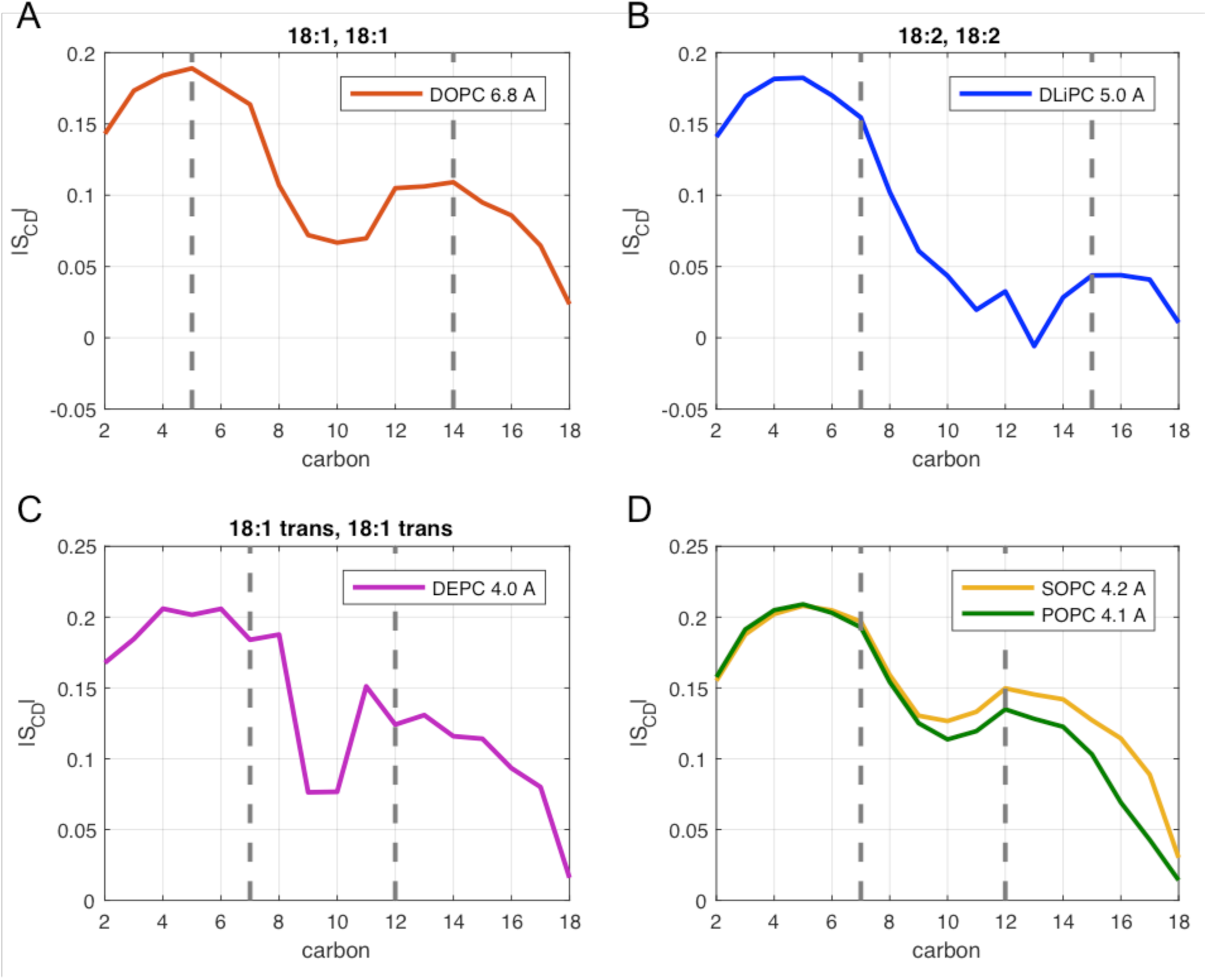
Heuristic approximation of the double bonds region (DB) in all unsaturated PC bilayers from Fig. 4. For lipids with two cis-unsaturated chains, DB extends 4 carbons above and below the midpoint of all double bonds; for bilayers with one saturated chain or two trans-unsaturated chains, DB extends 2 carbons above and below the midpoint of all double bonds. Shown are the acyl chain order parameter (*S*_CD_) profiles of the bilayers. Each profile is averaged over the two acyl chains of the lipids. Dashed lines surround the approximated double bonds region.

**Figure S8.**
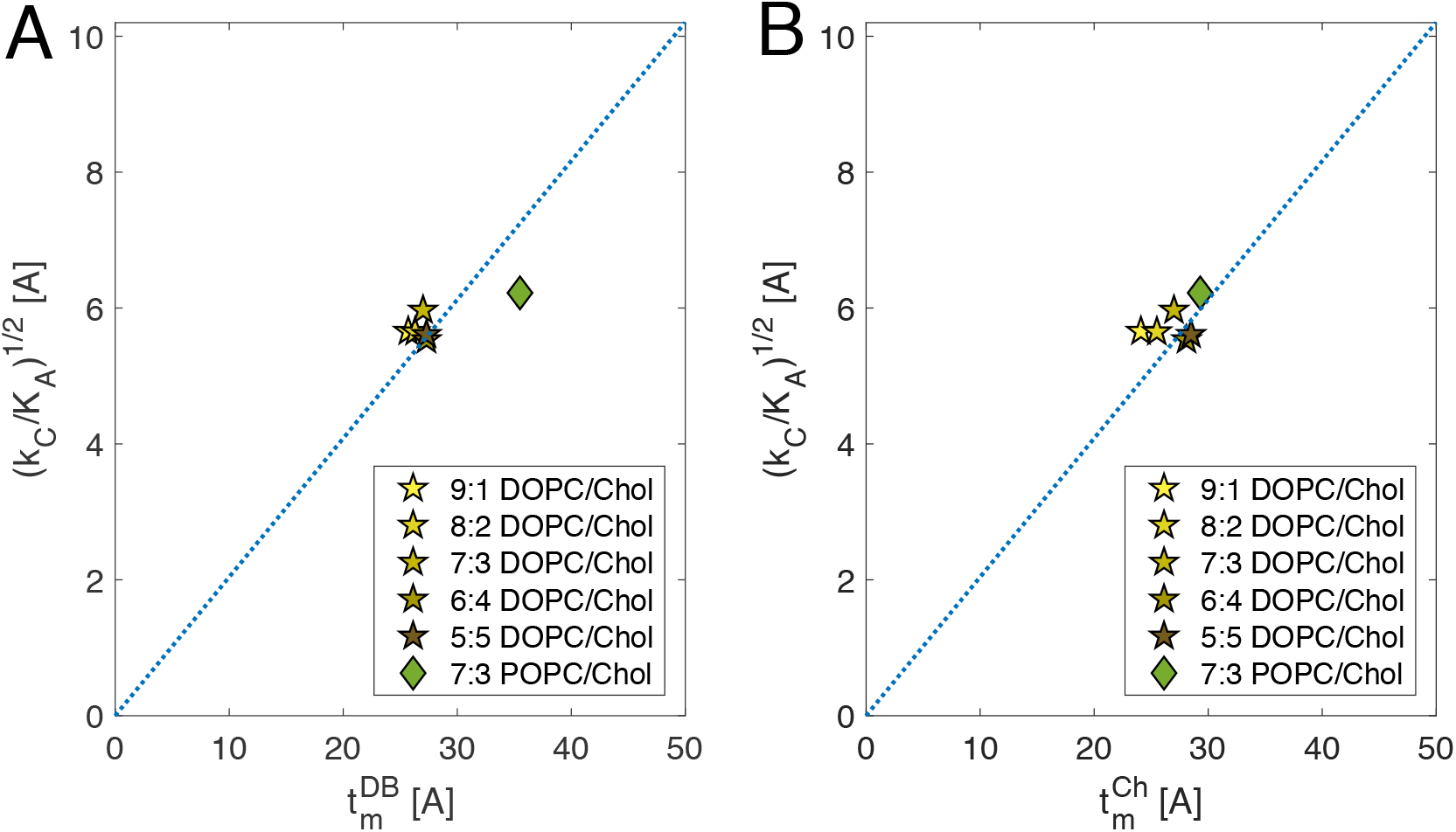
Application of the Polymer Brush Model (Eq. 17) to unsaturated bilayers with cholesterol using different definitions of *t_m_*: the phosphate-to-phosphate distance excluding either (in A) the perturbed region around the double bonds 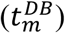, or (in B) the length of Chol’s ring body 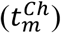. See Table S3 for exact quantities and discussion in the main text for more details.

**Figure S9.**
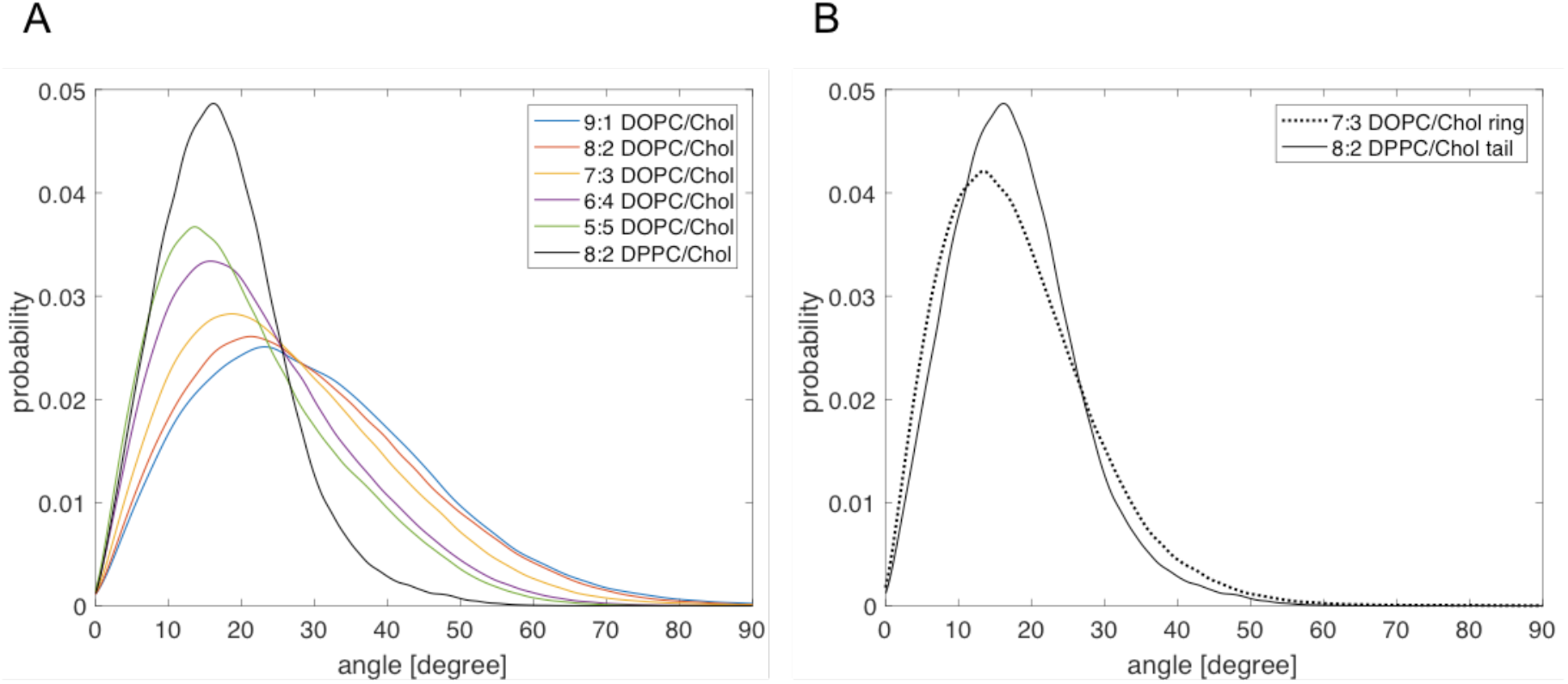
(**A**) Distribution of tilt angles of cholesterol’s tail (director vector connecting carbons 17 and 25 in CHARMM36 notation) with respect to bilayer normal (taken as the z dimension of the simulation box) for indicated cholesterol bilayers calculated from the simulation trajectories. Note that the distribution of Chol’s tail tilt angles for the DPPC/Chol bilayer is narrower than in the other bilayers. It is comparable to the distribution of Chol’s ring tilt angles shown in panel (**B**) (director vector connecting carbons 3 and 17 in CHARMM36 notation).

### Method Sections

#### S.1 Inter-block correlations and extended theoretical model

In order to evaluate the magnitude of the *Q* term appearing in Eq. 8 of the main text, we calculated the sum of the covariances of local areas in a leaflet, taking advantage of the relationship 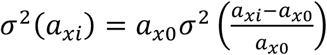. Since the relative changes in area can be expressed as characteristic changes in thickness (Eq. 11 of the main text), 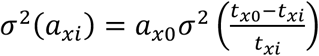 where *t* is thickness. Following the same approach as the one outlined in the “Leaflet compressibility from thickness fluctuations” section in the main text, we calculated the characteristic changes in the *relevant* leaflet thickness (see “Identifying the relevant thickness for fluctuations analysis” section in the main text) on an 8×8 Å^2^ grid in the leaflet. From that, we obtained the sum of the covariances of local areas as 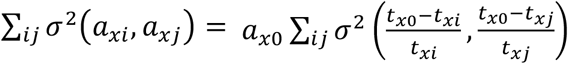. We then performed the 2D-bootstrapping algorithm outlined in Section S.7 to get the corresponding 95% confidence interval (CI). For three different bilayers representing very low (DPPC/Chol), medium (DPPC) and high (DOPC) fluidity, the 95% CI-s were [-0.002; 0.003], [-0.012; 0.018] and [-0.010; 0.014], respectively. Since all intervals contained 0, we can conclude that, empirically, the sum of the covariances of local areas within a leaflet is approximately 0 and consequently, *Q* ≈ 0.

The derivation of Eq. 10 in the main text (in particular, the transition from Eq. 7 to Eq. 8) assumes that the two leaflets are composed of the same number of elastic blocks. Since the smallest physically meaningful average unit area of a block is the average area of a lipid, and in an asymmetric membrane the two leaflets may have different number of lipids, in the following we extend the formulation to allow for different number of blocks in the two leaflets.

Let *n_x_* and *n_y_* be the number of blocks in the two leaflets such that *n_x_* ≠ *n_y_*. Since all blocks within a leaflet have the same compressibility modulus, i.e. the same variance, Eq. 7 can be written as:

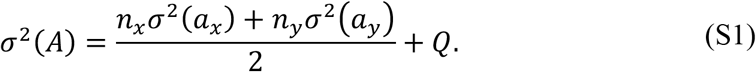

Further, since 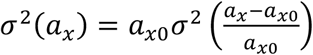,

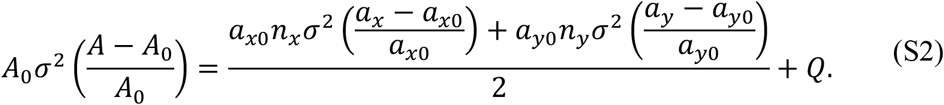

However, since *a_x0_n_x_* = *a_y0_n_y_* = *A_0_* and 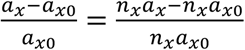, Eq. S2 simplifies to:

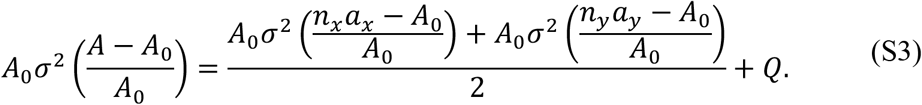

Since 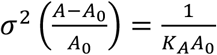, the above leads to:

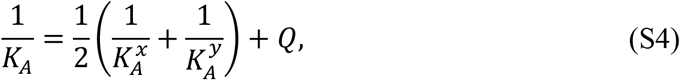

which is the same expression as Eq. 8.

#### S.2 Two approaches for calculating *K_A_*

As mentioned in the main text, 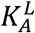 can be obtained either from the equipartition theorem (Eq. 12) or from statistical mechanics (Eq. 13). Eq. 12 relies on an accurate estimation of the mean of the squared characteristic changes in thickness 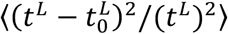. However, this calculation becomes problematic because of numerical issues. We illustrate the problem in Fig. S10 by using as an example the local relevant thicknesses calculated for the top leaflet of DPPC as discussed in the main text. Fig. S10A shows the distribution of the *relative* changes in thickness while Fig. S10B shows the corresponding distribution of *characteristic* changes in thickness. The latter is more skewed and has a long tail to the right (beyond the x-axis limits on the plot) coming from outliers, e.g. very small local thicknesses 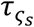 that produce very large characteristic changes since 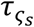 appears in the denominator. Once squared, the distribution becomes even more skewed and sensitive to outliers as illustrated in Fig. S10C (it again has a very long tail that is truncated in the figure). The accurate estimation of the mean from the distribution in Fig. S10C thus becomes challenging. Recognizing the fact that theoretically Eqs. 12–13 are valid only in the regime of small deformations around the mean thickness, and that 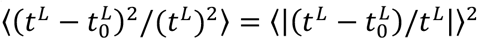, we calculated 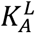 from Eq. 12 by considering only *t^L^* within some *p* percent of 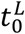. As can be seen in the examples in Fig. S11, the results depend strongly on *p*.

**Figure S10.**
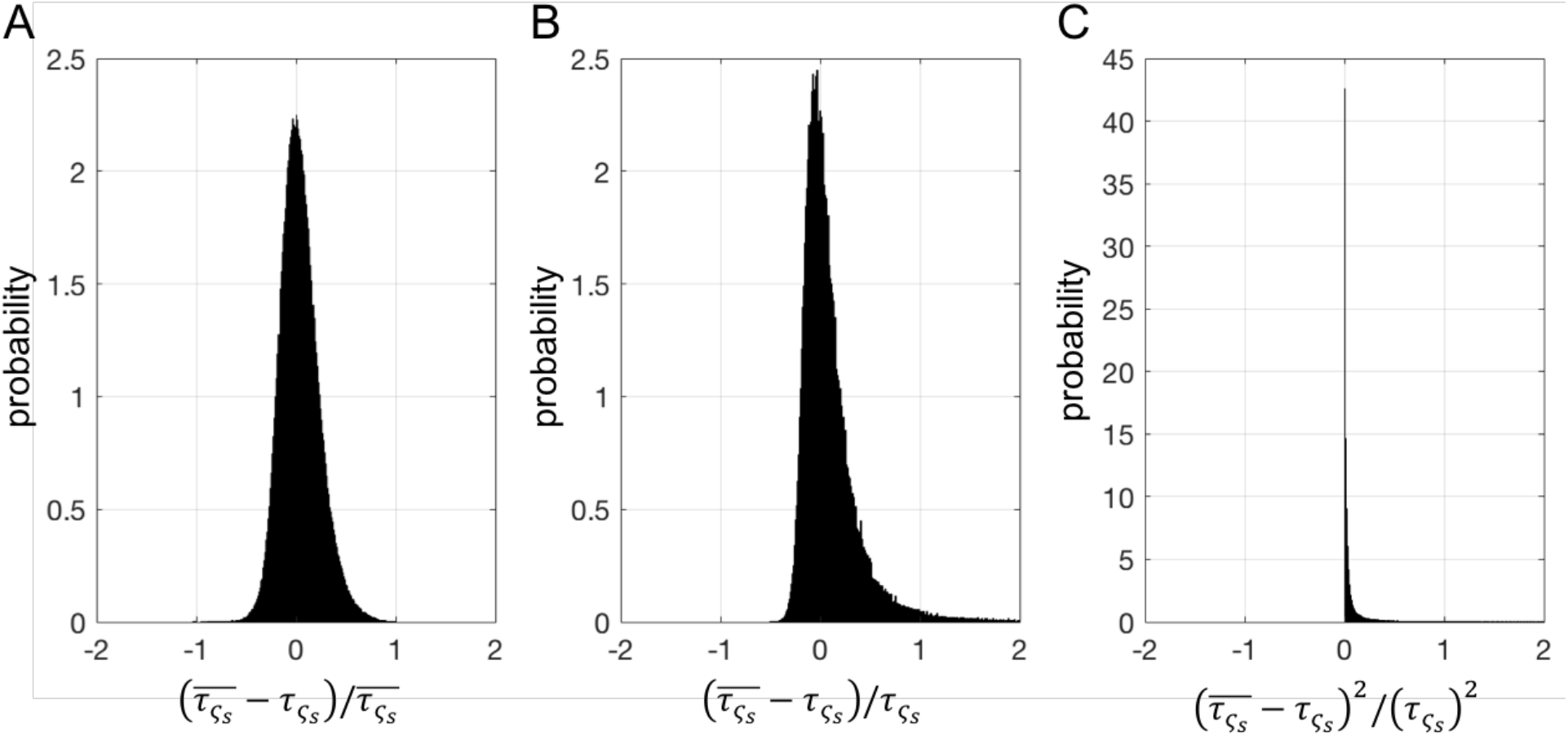
Probability density functions of different changes in relevant thickness for the top leaflet of DPPC: relative (A), characteristic (B) and squared characteristic (C) changes in thickness.

In contrast, obtaining 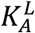 from Eq. 13 requires an accurate representation of the *non-squared* distribution within a small region around 0. Furthermore, Taylor expansion of the expression for the energy (Eq. 11) shows that the distributions of relative and characteristic changes in thickness are the same for thicknesses around the mean 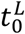, i.e.:

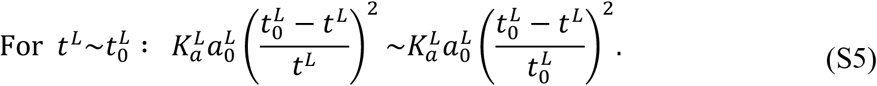

**Figure S11.**
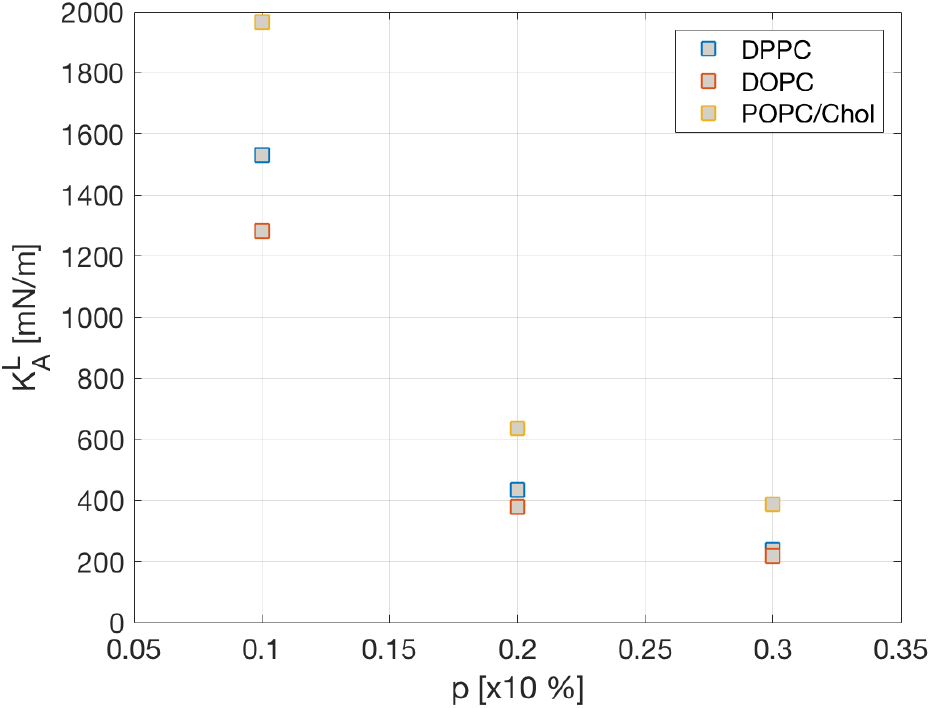
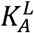 calculated with Eq. 12 in the main text by using only thicknesses within *p* percent of the mean thickness 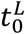. Results are shown for the top leaflets of DPPC (blue), DOPC (red) and POPC/Chol (yellow) bilayers.

This allows us to use the distribution of relative thicknesses (Fig. S10A) to obtain 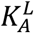 without running into issues arising from outliers in the local thicknesses (note that small local thicknesses result in relative changes ~1 that do not affect the PMF analysis which is done on thicknesses with relative changes within the range −0.07 and 0.07).

#### S.3 Averaging lipid chains for different lipids and lipid mixtures

Eq. 15 can be applied directly to single component bilayers with lipids that have multiple chains of the same length such as DPPC, DOPC, DMPC, or TOCL (with the special treatment of double bonds as described in the text). For lipids that have different length chains we use the following algorithm. If *N*_1_ and *N*_2_ are the lengths of chains 1 and 2 of a two-tailed lipid (i.e. the number of carbon atoms in the chain after any averaging for the double bonds), and *N*_1_ < *N*_2_, then:

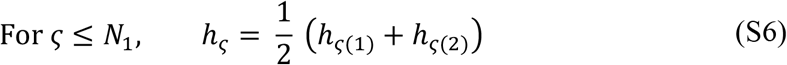

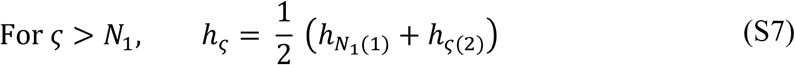

In other words, for carbons 1 through *N*_1_ we average the heights of the two chains. For carbons on chain 2 greater than *N*_2_, we average the height of the carbon with the height of the terminal methyl carbon of chain 1.

For PSM, the first carbons not attached to oxygen on the two chains are C4S and C2F (using CHARMM36 atom name notation). Hence, both chains have the same length and their heights are averaged using Eq. 15.

For lipid mixtures, we do not perform the interpolation separately for the different lipid components but instead treat all of them the same way: Assuming that all lipids have the same number of chains (e.g. 2 as in most cases), the height of the C carbon of chain CH is calculated from the surface constructed from the C carbons of chain CH on all lipids in the leaflet. If chain CH of one of the lipids (lipid X) is shorter than the CH chains of the other lipids, for the interpolated surfaces of subsequent carbons further down the chain, we use the methyl carbon of the CH chain of lipid X. This ensures that the interpolated surface always contains exactly 1 atom from each lipid molecule irrespective of the lipid type. This leads to interpolated surfaces for the carbons of two chains (if one of the chains is unsaturated for all lipids in the mixture, we apply the same treatment to the double bonds as in the single component unsaturated bilayers) and we proceed with the averaging of their heights as described above.

If cholesterol is present in a leaflet, we exclude it from the calculation of the interpolated heights. That is, the interpolation is performed only on the non-cholesterol components. Therefore, in order to calculate 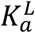 with Eq. 13 we need to find the average area per non-Chol molecules 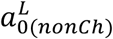 To this end, we use the simple model proposed by Alwarawrah et al. whereby the partial area of Chol, 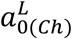, is approximated from Chol’s average tilt angle (Eq. 8 in Ref [7]). 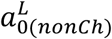 is then obtained from the relationship:

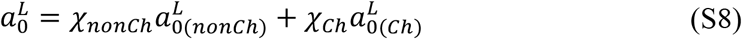

where *χ_Ch_* is Chol’s mole fraction in the leaflet and *χ_nonCh_ =* 1−*χ_Ch_* is the mole fraction of the non-Chol components. Note that the influence of cholesterol on the membrane compressibility is still considered in the formulation implicitly through its effect on the local thickness/area fluctuations of the membrane surface (see Simulations analysis and method implementation section in the text for more details, including a necessary correction applied for high mole fractions of Chol).

#### S.4 Determining the effective area of normalization

In Eq. 14, *ς* is unique for a lipid. The summations are performed over all lipids in the leaflet with the contribution of each lipid being weighted by 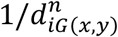 and thus decreasing the further away the lipid’s atom *ς* is from the grid point. Therefore, the *effective* area over which the surface heights *h_ς_* and consequently, thicknesses *τ_ς_*, are calculated depends on the interpolation order *n*: The higher the interpolation order, the more local the analysis, i.e. the higher the relative contribution of the atoms closest to the grid point. The effective area can thus be approximated by 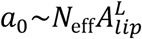 where 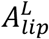 is the equilibrium area per lipid in the leaflet, and 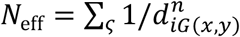 is the sum of all weights that represents the *effective* number of lipids contributing to *h*_ς,*G*(*x*,*y*)_. For example, if *n* = 0 then *N*_eff_ = ∑_ς_ 1/1 = *N* where *N* is the total number of lipids in the leaflet, and 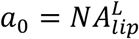. Stated differently, with 0^th^ order interpolation the heights at all grid points are the same and equal to the average z position across all atoms of type ς in the leaflet. For *n* > 1,

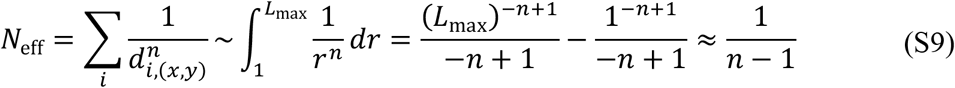

where the second equality comes from the power rule, *r* is a variable of integration representative of distance, and 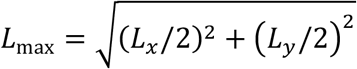 is the maximum 2D distance between two points on the leaflet surface, given periodic boundary conditions (*L_x_* and *L_y_* are the lateral dimensions of the simulation box). If *a*_0,*n*_ denotes the *effective* area in the interpolation scheme with interpolation order *n*, then from Eq. S9 it follows that:

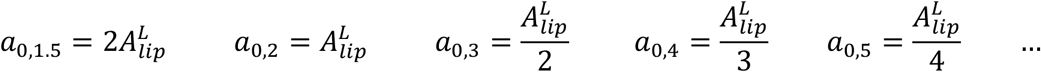

Note that *n* = 1 is a special case where the integral in Eq. S9 can be approximated with ln *L*_max_ and 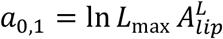. Fig. S2 shows a comparison between the *effective* compressibilities calculated with different *n* at different carbons. Since *a*_0,2_ is equal to the equilibrium area per lipid,for convenience we choose *n* = 2 and perform all subsequent analysis using the equation:

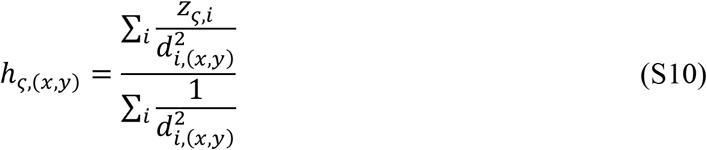

#### S.5 Testing the volume incompressibility assumption

The conversion from changes in area to changes in thickness in Eq. 11 of the main text relies on the assumption of volume incompressibility that enforces the preservation of the product of area and thickness. In our model this assumption is applied at the level of individual lipid molecules (i.e. the elastic blocks that make up a leaflet). To test its validity, we first took one of the trajectories of a bilayer with a relatively loose packing density and examined how the lipid volume varied in the course of the simulation. To calculate the volume of each lipid, we performed a 3-dimensional Voronoi analysis in which the space in every trajectory frame was partitioned so that every atom was assigned a 3D Voronoi voxel (taking into account periodic boundary conditions). The volume of each lipid was then obtained from the sum of the voxel volumes of all lipid atoms. As shown in Fig. S12, the resulting volume for DOPC had a mean of 1312 Å3, which is very similar to the experimentally measured value of 1295 Å3 [8], and a standard deviation of 33.4 Å^3^ (or 2.5% of the mean). Thus, while not constant, the volume exhibited only a very small variation around the mean.

**Figure S12.**
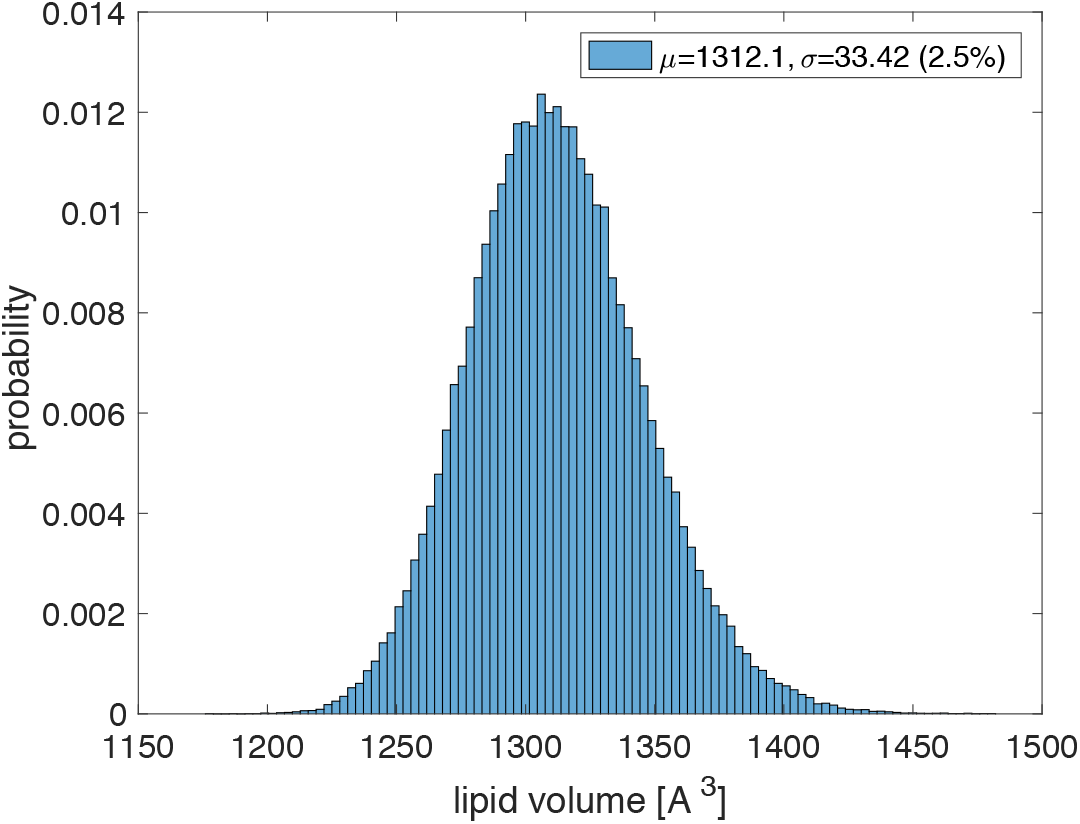
Probability distribution of the lipid volume of DOPC. Lipid volume was calculated from 3D Voronoi analysis as explained in the text. The histogram was constructed from the time-resolved volumes of all DOPC lipids in the bilayer.

To evaluate the effect of small variations in the volume on the calculated area compressibility modulus, we expressed the volume *V* as a normalized normal distribution described by the parameters *V*_0_ (its mean) and *σ* (its standard deviation):

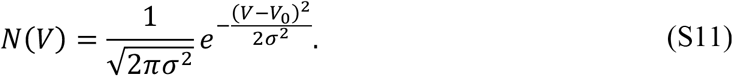

Now consider the function:

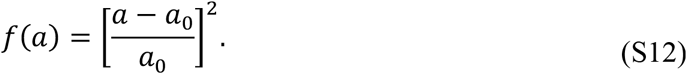

The expectation value of *f* over the normal distribution *N*(*V*) is:

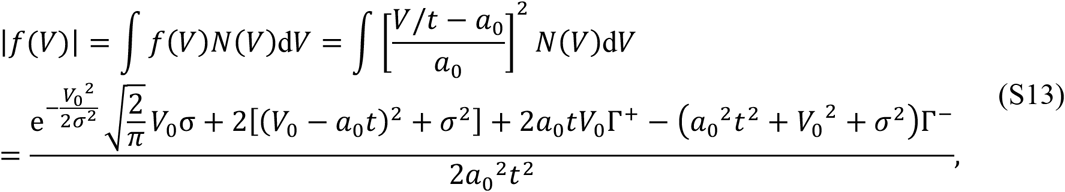

where *a*_0_ = *V*_0_/*t*_0_ and Γ^+^ and Γ^−^ are defined as:

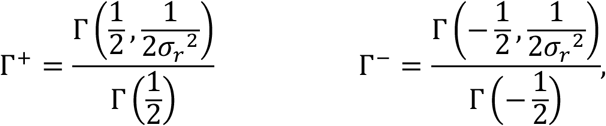

with *σ_r_* = *σ*/*V*_3_ being the relative change in the volume, and the *Gamma* and *incomplete Gamma* functions, Γ(*z*) and Γ(*a*,*z*) respectively, defined as:

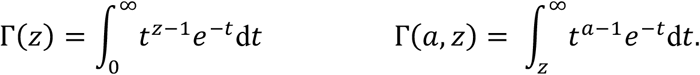

Thus, expressed in terms of *σ_r_* and the change in thickness 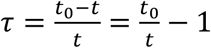 becomes:

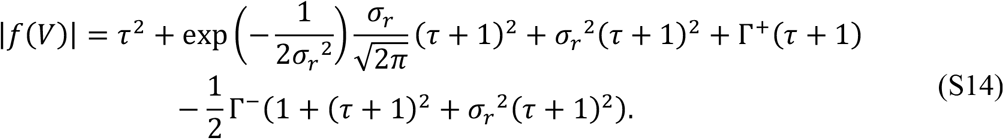

If *V* is a constant, i.e. as *σ_r_* → 0, |*f* (*V*)| = *τ*^2^. Fig. S13A shows |(*V*)| as a function of *τ* for the cases when *σ_r_* → 0, i.e. when *V* is constant, and *σ_r_* = 0.025, i.e. when *V* varies by 2.5% as in Fig. S12. The two cases produce very similar profiles, indicating that the differences between them are, if anything, very small. Fig. S13B further shows the relative changes and corresponding error of *K_A_* as a function of *σ_r_*. As evident from the plots, for changes in the volume of up to 4% (*σ_r_* = 0.04), the error of *K_A_* is less than 1%. Since this is well within the error of the calculated compressibility moduli (see Tables 1 and 2 in the main text), we conclude that the volume incompressibility assumption is a reasonable approximation at the level of individual lipid molecules and can thus be used safely in our framework.

**Figure S13.**
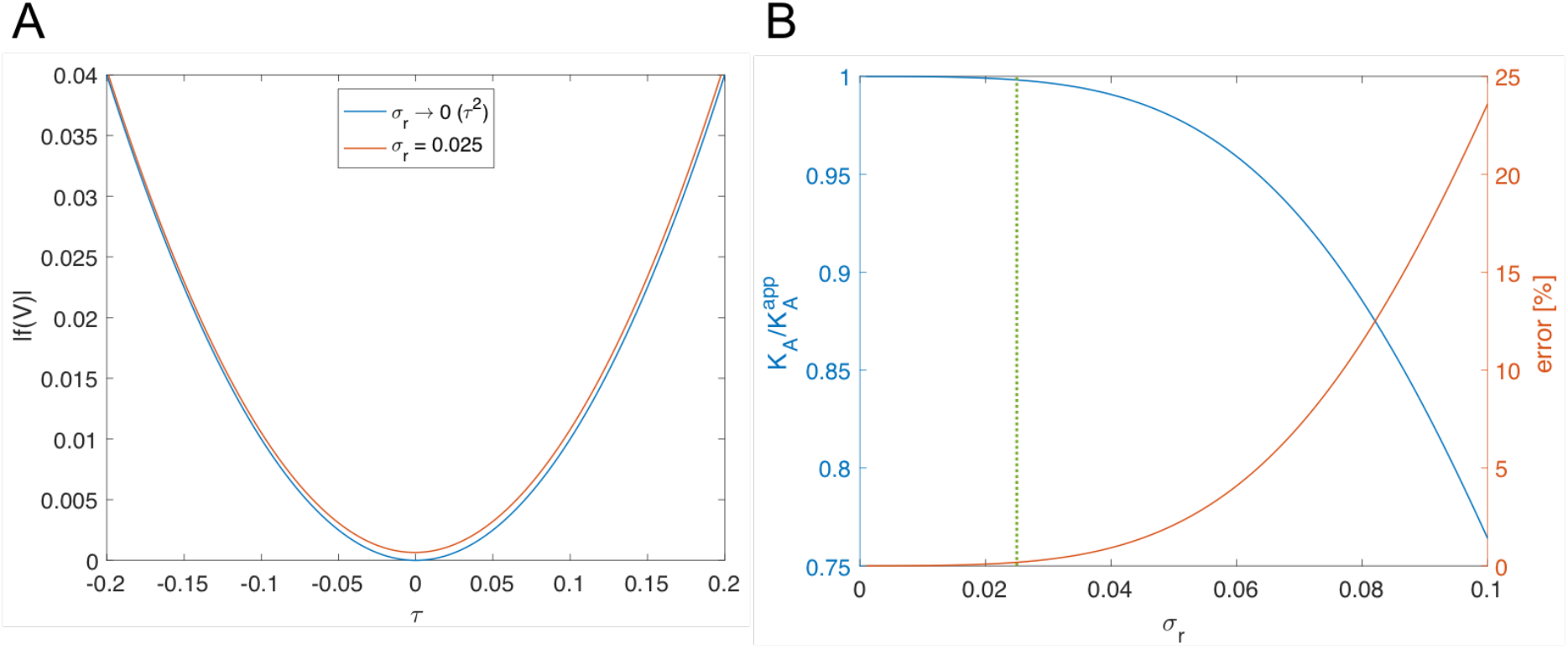
**A.** The expectation value of *f* over the normal distribution *N*(V) for relative change in the volume of 0% (blue) and 2.5% (red). **B.** Relative changes in *K_A_* (blue, left) and corresponding errors (red, right) for different values of *σ_r_* calculated by fitting a quadratic function to |*f*(V)| in the region −0.07 ≤ τ ≤ 0.07 (see Section S.6). For changes in the volume of up to 4% (*σ_r_* = 0.04), the error of *K_A_* is less than 1%.

#### S.6 Calculating 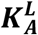 from the relevant local thicknesses

Once the relevant thickness has been identified and the corresponding PMF is estimated (see Fig. 2B in the main text), we fit a quadratic function to the PMF in a small region around the mean thickness in order to recover 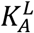. As mentioned above (see Fig. S10 and Eq. S5), for small thicknesses around the mean thickness the distributions of relative and characteristic changes are the same. For practical reasons (see above) we use the relative thickness changes for the PMF fitting. In particular, from the kernel density of the thicknesses (Fig. 2A), we estimate the PMF of the relative changes in thickness (similar to the left-hand-side of Eq. 13). To identify the region for fitting, we search within a small range of thicknesses between 5 and 7% of the mean thickness. This range ensures that the analyzed thickness deformations are small enough to satisfy the theoretical requirements of the model (i.e. quadratic approximation of the energy), while at the same time providing sufficient sampling of the data, minimally affected by noise features in the underlying distribution. The full protocol goes as follows:

1. Estimate the thickness distribution with a kernel density estimation using the *ksdensity* function in MATLAB. The bandwidth, *bw* (i.e. smoothing parameter) of the kernel density is determined automatically based on the data. From the kernel density, find 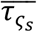 corresponding to the peak of the kernel distribution.
2. Use the kernel weights to estimate the probability distribution of the relative thickness changes, 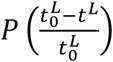, and the corresponding 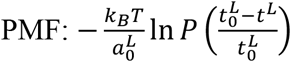.
3. Select a range of relative thickness changes (between 0.05 and 0.07), corresponding to changes in thickness within 5 to 7% of the mean thickness.
4. Sample from the estimated probability distribution within the thickness range (i.e. sample the relative thickness changes (raw data) within the selected range and add Gaussian noise to them with bandwidth equal to 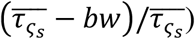.
5. Construct a qqplot of the sampled data, i.e. compare the quantiles of the sampled distribution to those of a normal distribution. Calculate the fraction, *fracq* of the sampled data quantilies that are within 0.01 of the normal distribution quantiles. The larger *fracq*, the closer the sampled distribution is to a normal distribution.
6. For each range (in step 3), repeat steps 4-5 ten times and find the mean *fracq*. The range with maximum mean *fracq* (i.e. closest to a normal distribution) is the range that we choose for subsequent PMF fitting.
7. Fit a quadratic function of the form *f*(*x*) = *c*_1_*x*^2^+*c*_2_ to the PMF in the identified region. 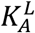 is the coefficient of the quadratic term (*c*_1_) in the best fit.

#### S.7 Error on leaflet compressibility

The calculation of 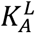 from local changes in thickness is based on sampling both in time (*N* time points over the course of the trajectory) and space (*G_x_G_y_* grid points across the two dimensional grid on the leaflet surface). However, there can be correlations both in time (slow motions leading to autocorrelation) and in space (local correlated motions due to interaction between atoms), and thus the effective number of independent observations of the leaflet thickness may be less than *N G_x_G_y_*. In order to account for these correlations in our estimate of the error on 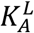, we use a 2-dimensional non-parametric moving block bootstrapping approach in which we resample blocks of data in both time and space. The procedure goes as follows:

1. We first find for each height *h* the median autocorrelation time *ξ_h_* across all grid points. The largest *ξ_h_* determines the number of consecutive time points, *N_SEG_*, in a resampled time block.
2. We then find the maximum number of grid points *G_SEG_* such that 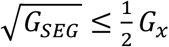 and 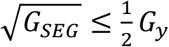.
3. We resample a block of *N_SEG_ G_SEG_* data points by picking a random point in time (*T_p_*) between 1 and *N*−*N_SEG_*, and space (*S_p_*), and taking *N_SEG_* consecutive frames starting from *T_p_* and a square patch of *G_SEG_* grid points with a top left corner at *S_p_* (using periodic boundary conditions).
4. We repeat step 3 *η* times where *η* = max (*N*/*N_SEG_ G_SEG_*) is an integer, to construct a resampled data set *D*′ with approximately the same size as the original data set, i.e. *ηN_SEG_ G_SEG_* ≈ *N G_x_ G_y_*. We calculate the compressibility modulus from *D*′, i.e. 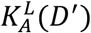.
5. We repeat step 4 one hundred times and obtain the error of 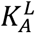 from the standard deviation of the set of 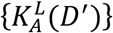 values.

#### S.8 NPγT simulations

For some of the bilayers, a series of simulations at fixed non-zero tension were performed in order to obtain the bilayer *K_A_* directly from the relationship between applied tension and resulting area expansion, as done in micropipette experiments [9]. The choice of tensions to apply for each system (listed in Table S2) was based on a few different factors: We made sure to have at least one negative and two positive non-zero tensions, so that the linear fit for extracting *K_A_* is performed on at least 4 data points (including the 0-tension bilayer). Depending on the compressibility of the bilayer, the exact γ values for the system were also chosen to produce significantly different relative changes in the area but without being too large due to deviation from linearity at higher tensions.

